# Synaptic plasticity regulated by phosphorylation of PSD-95 Serine 73 in dorsal CA1 is required for contextual fear extinction

**DOI:** 10.1101/2020.11.13.381368

**Authors:** Magdalena Ziółkowska, Malgorzata Borczyk, Agata Nowacka, Maria Nalberczak-Skóra, Małgorzata Alicja Śliwińska, Magdalena Robacha, Kacper Łukasiewicz, Anna Cały, Edyta Skonieczna, Kamil F. Tomaszewski, Tomasz Wójtowicz, Jakub Włodarczyk, Tytus Bernaś, Ahmad Salamian, Kasia Radwanska

## Abstract

The ability to extinguish fearful memories is essential for survival. Accumulating data indicate that the dorsal CA1 area (dCA1) contributes to this process. However, the cellular and molecular basis of fear memory extinction remains poorly understood. Postsynaptic density protein 95 (PSD-95) regulates the structure and function of glutamatergic synapses. Here, using dCA1-targeted genetic and chemogenetic manipulations *in vivo* combined with PSD-95 immunostaining and 3D electron microscopy *ex vivo*, we demonstrate that phosphorylation of PSD-95 at serine 73 PSD-95(S73) is necessary for contextual fear extinction-induced expression of PSD-95 and synaptic plasticity. Moreover, PSD-95(S73) phosphorylation is not necessary for fear memory formation and recall but is required for extinction of contextual fear. Overall, our data shows how PSD-95-dependent synaptic plasticity in the hippocampus contributes to the persistence of fear memories.

## INTRODUCTION

The ability to form, store, and update fearful memories is essential for animal survival. In mammals, the formation and updating of such memories involve the hippocampus (Baldi and Bucherelli, 2015; Frankland and Bontempi, 2005; Neves et al., 2008). In particular, the formation of contextual fear memories strengthens Schaffer collaterals in the dorsal CA1 area (dCA1) through NMDA receptor-dependent Hebbian forms of synaptic plasticity (Abraham et al., 2019; Bliss and Collingridge, 1993; Morris et al., 2003) linked with growth and addition of new dendritic spines (harboring glutamatergic synapses) (Aziz et al., 2019; Mahmmoud et al., 2015; Radwanska et al., 2011; Restivo et al., 2009). Similarly, contextual fear extinction induces functional, structural, and molecular alterations of dCA1 synapses (Garín-Aguilar et al., 2012; Schuette et al., 2020; Stansley et al., 2018). While the role of dCA1 synaptic plasticity in contextual fear memory formation has been recently questioned (Bannerman et al., 2014, 2012), its role in contextual fear memory extinction is mostly unknown. Understanding the molecular and cellular mechanisms that underlie fear extinction memory is crucial to develop new therapeutic approaches to alleviate persistent and unmalleable fear.

PSD-95 is the major scaffolding protein of a glutamatergic synapse (Cheng et al., 2006), affecting its stability and maturation (Ehrlich et al., 2007; Steiner et al., 2008; Sturgill et al., 2009; Taft and Turrigiano, 2014) as well as functional (Béïque and Andrade, 2003; Ehrlich and Malinow, 2004; Migaud et al., 1998; Stein et al., 2003) and structural plasticity (Chen et al., 2011; Nikonenko et al., 2008; Steiner et al., 2008). PSD-95 interacts directly with NMDA receptors and through an auxiliary protein stargazin with AMPA receptors (Kornau et al., 1995; Schnell et al., 2002). Interaction of PSD-95 with stargazin regulates the synaptic content of AMPARs (Bats et al., 2007; Chetkovich et al., 2002; Schnell et al., 2002). In agreement with these findings, overexpression of PSD-95 occludes long-term potentiation (LTP) (Ehrlich and Malinow, 2004; Stein et al., 2003) and decreases the threshold for long-term depression (LTD) induction (Béïque and Andrade, 2003). Conversely, mice lacking functional PSD-95 protein have greatly enhanced hippocampal, NMDAR-dependent LTP, whereas NMDAR-dependent LTD is absent (Migaud et al., 1998). Synaptic localisation of PSD-95 is controlled by a range of posttranslational modifications with opposing effects on synaptic retention (Vallejo et al., 2017). One such modification is the phosphorylation of Serine 73 (S73). It was first described as a target of αCaMKII that promotes PSD-95 dissociation from the NMDA receptor subunit NR2A (Gardoni et al., 2006). Further studies showed that S73 phosphorylation induces PSD-95 activity-dependent trafficking that is necessary for termination of synaptic growth after NMDAR stimulation, as well as PSD-95 downregulation during NMDAR-dependent LTD (Nowacka et al., 2020; Steiner et al., 2008). Interestingly, the loss-of-function PSD-95 mutant mice lacking the guanylate kinase domain of PSD-95 (Migaud et al., 1998) show normal contextual fear memory but impaired extinction of contextual fear (Fitzgerald et al., 2015), indicating that PSD-95-dependent synaptic plasticity contributes to the updating rather than the formation of contextual fear memory. The function of PSD-95(S73), or other PSD-95 modifications, in memory processes is mostly unknown. Here, we hypothesized that PSD-95(S73)-dependent synaptic plasticity in dCA1 contributes to extinction of contextual fear memories.

The present study tests this hypothesis by integrated analyses of PSD-95 protein expression and dendritic spines morphology with nanoscale resolution, combined with genetic and chemogenetic manipulations and behavioral studies. Using dCA1-targeted overexpression of PSD-95 and chemogenetic manipulations, we show that phosphorylation of PSD-95(S73) is necessary for contextual fear extinction-induced PSD-95 expression and remodeling of dendritic spines. Moreover, it is not necessary for fear memory formation but required for fear extinction even after extensive fear extinction training. Overall, our data indicate that the dCA1 PSD-95(S73)-driven synaptic processes during the extinction of fear memories enable extinction of the contextual fear memory.

## RESULTS

Previous data indicate that loss-of-function mutant mice lacking the guanylate kinase domain of PSD-95 do not show contextual fear extinction, while contextual memory formation is intact (Fitzgerald et al., 2015). Moreover, the contextual fear extinction induces dendritic spine remodeling in the dorsal CA1 area (dCA1) (Garín-Aguilar et al., 2012). Based on these findings, we hypothesise that PSD-95 controls extinction-induced remodeling of dCA1 neuronal circuits supporting contextual fear memory extinction.

### Acquisition and extinction of contextual fear memory

To study the synaptic mechanisms of contextual fear extinction memory, we used Pavlovian contextual fear conditioning. Mice were exposed to a new context, and 5 electric shocks (5US) were delivered. The fear memory was extinguished the next day by re-exposure to the same context without the delivery of USs (Figure S1). Mice showed low freezing levels in a novel context before delivery of electric shocks (pre-US), and freezing increased during the training (post-US), indicating fear memory formation. Twenty-four hours later, mice were re-exposed for 30 minutes to the training context without the US’s presentation for the fear extinction memory session (Extinction). Freezing levels were high at the beginning of the session, indicating fear memory retrieval and decreased within the session, indicating extinction learning. Twenty-four hours later, we tested for 5 minutes the consolidation of fear extinction memory (Test). During the Test, freezing levels were lower than at the beginning of Extinction, indicating long-term fear extinction memory formation.

### The effect of contextual fear extinction memory on PSD-95 expression in dCA1

To investigate the role of PSD-95 in contextual fear memory consolidation and extinction, we analysed the expression of PSD-95 protein in Thy1-GFP(M) mice (Feng et al., 2000). Thy1-GFP(M) mice express GFP in a sparsely distributed population of the glutamatergic neurons, allowing for dendritic spines visualisation (Figure 1A). The expression of PSD-95 protein, and its colocalization with dendritic protrusions, were analysed in three domains of dCA1: stratum oriens (stOri), stratum radiatum (stRad) and stratum lacunosum-moleculare (stLM) (Figure 1B-C). We analysed these regions separately as previous data found dendrite-specific long-term dendritic spines changes after contextual fear conditioning (Restivo et al., 2009).

**Figure 1.**
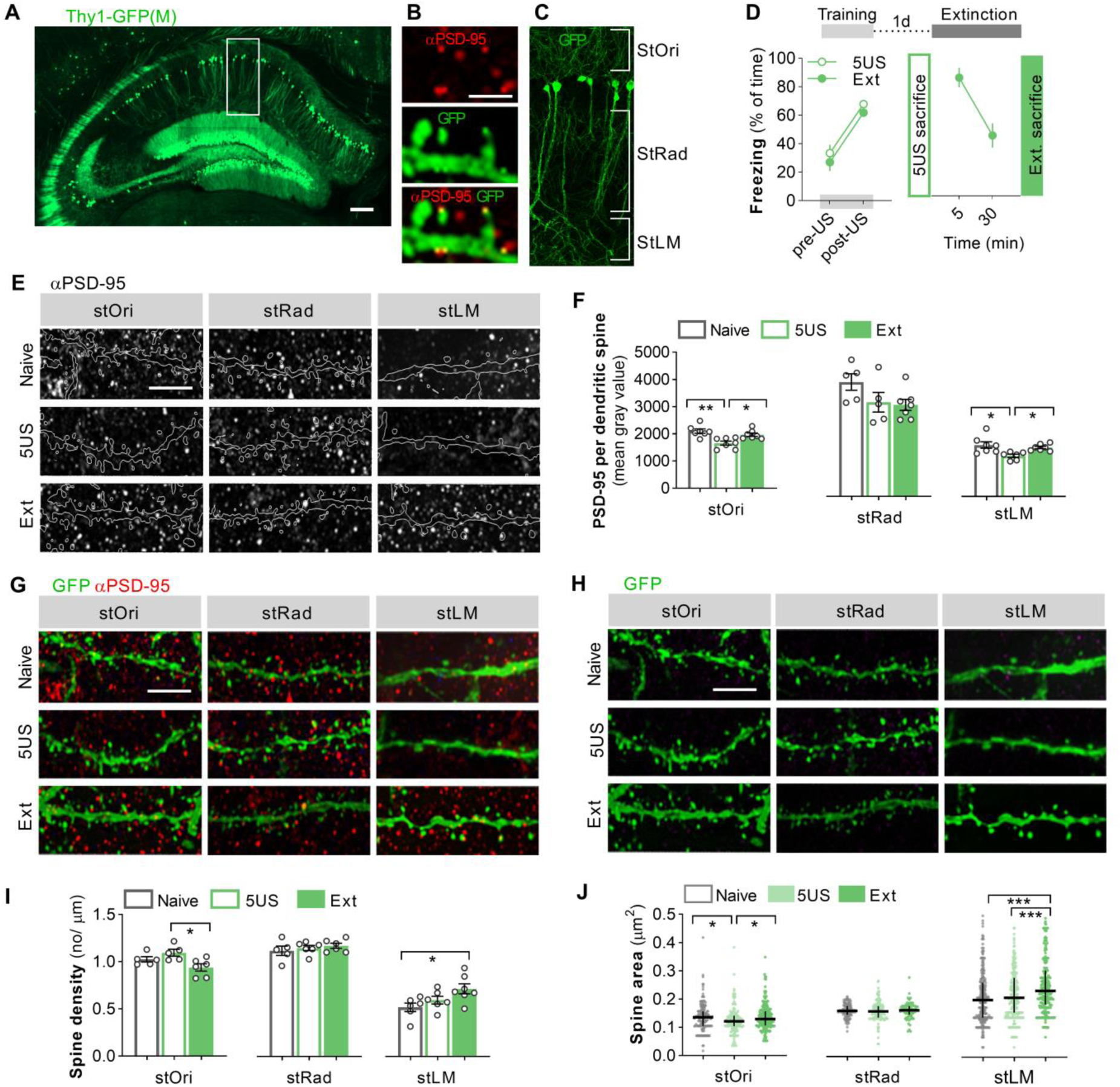
Formation and extinction of contextual fear memory regulate expression of synaptic PSD-95 protein and remodeling of dendritic spines in dCA1. **(A-C)** Dendritic spines were analysed in three domains of the dendritic tree of dCA1 pyramidal neurons in Thy1-GFP(M) mice: stOri, stRad and stLM. **(A)** Microphotography of dCA1 of a Thy1-GFP(M) mouse. **(B)** High magnification of confocal scans showing colocalization of PSD-95 immunostaining and dendritic spines. **(C)** Division of dCA1 dendritic tree domains. **(D)** Experimental timeline and freezing levels of mice from two experimental groups: fear conditioning training (5US, n = 6) only and fear extinction (Ext, n = 7). **(E)** Representative confocal images of PSD-95 immunostaining (maximum projections of z-stacks composed of 20 scans) are shown for three domains of the dendritic tree. (**F**) Summary of data showing PSD-95 expression in stOri (mouse/spine: Naïve = 6/579; 5US = 6/807; Ext = 7/986), stRad (mouse/spine: Naïve = 6/571; 5US = 6/619; Ext = 7/712), and stLM (mouse/spine: Naïve = 6/705; 5US = 6/650; Ext = 7/925). **(G-H)** Representative confocal images of dendrites colocalized with PSD-95 immunostaining from Thy1-GFP(M) mice that underwent training are shown for three domains of the dendritic tree. **(I)** Summary of data showing dendritic spines density in stOri (mouse/dendrite: Naïve = 6/16; 5US = 6/24; Ext = 7/34), stRad (mouse/dendrite: Naïve = 6/18; 5US = 6/20; Ext = 7/19), and stLM (mice/dendrite: Naïve = 6/31; 5US = 6/25; Ext = 7/37). **(J)** Summary of data showing average dendritic spine area in stOri, (mice/ spines: Naïve = 6/579; 5US = 6/807; Ext = 7/986), stRad (mouse/spine: Naïve = 6/571; 5US = 6/619; Ext = 7/712), and stLM (mouse/spine Naïve = 6/705; 5US = 6/650; Ext = 7/925). For F and I, each dot represents one mouse. For J, each dot represents one dendritic spine. Scale bars: A: 0.5 mm, B: 8 μm, E, G, H: 15 μm. *P < 0.05, **P < 0.01; ***P < 0.001.

Thy1-GFP(M) mice underwent contextual fear conditioning. They showed low freezing levels in the novel context before delivery of electric shocks, after which freezing levels increased the remainder of the training session (Figure 1D) (RM ANOVA, effect of time: F(1, 7) = 734.1, P < 0.0001). Twenty-four hours later, one group of mice was sacrificed (5US), and the second group was re-exposed to the training context without presentation of US for fear extinction (Ext). Freezing levels were high at the beginning of the session and decreased within the session (Figure 1D) (t = 3.720, df = 6, P < 0.001). Mice were sacrificed immediately after the fear extinction session. As controls, naïve mice were taken from their home cages. The analysis of PSD-95 immunostaining revealed a significant effect of training (RM ANOVA, F(2, 22) = 7.69, P = 0.003) and dCA1 region (F(1.317, 18.44) = 141.0; P < 0.001) on PSD-95 expression per dendritic spine (Figure 1F). *Post hoc* tests indicated that in the stOri and stLM, PSD-95 expression decreased in the 5US group, compared to the Naïve mice (Tukey’s multiple comparisons test, stOri: P = 0.004; stLM: P = 0.038), and increased after extinction (Ext), compared to 5US group (stOri: P = 0.019; stLM: P = 0.009) (Figure 1F). No difference in PSD-95 levels was observed in stRad between the groups. Thus, our data show that both the formation and extinction of contextual fear memory regulate PSD-95 levels in dCA1 strata, and the effect is specific to stOri and stLM regions.

Since PSD-95 is expressed in large and mature spines (El-Husseini et al., 2000), we checked whether the changes in PSD-95 levels were associated with dendritic spine remodelling. We did not observe a significant effect of training (RM ANOVA, F(2, 48) = 3.149, P = 0.052), but we did discover a region effect (F(1.788, 42.92) = 7.381, P = 0.002) and training × region interaction (F(4, 48) = 5.48, P = 0.001) on dendritic spines density (Figure 1I). In stOri, dendritic spines density decreased after fear extinction training (Ext) compared to the trained mice (5US) (Tukey’s test, P = 0.025) (Figure 1I). In stLM, dendritic spine density was increased in the Ext group compared to the Naïve mice (P = 0.039). No changes in spine density were observed in the stRad. Moreover, we found a significant effect of training on the median area of dendritic spines in the stOri (Kruskal-Wallis test, H = 8.921, P = 0.012) and stLM (H = 28.074, P < 0.001), but not stRad (H = 5.919, P = 0.744) (Figure 1J). In stOri, the median spine area was decreased in the 5US group compared to the Naïve mice (Dunn’s multiple comparisons test, P = 0.032) and increased after extinction (Ext) compared to the 5US group (P = 0.02). In stLM, the median spine area did not change after training (5US), compared to the Naïve mice (P > 0.05), but increased after extinction (Ext), compared to the 5US group (P = 0.005). Thus, increased expression of PSD-95 per dendritic spine in stOri and stLM during contextual fear extinction, as compared to the 5US group, was coupled with an increased median spine area. Overall, our data indicate remodelling of the dCA1 neuronal circuits during contextual fear extinction that presumably involves upregulation of PSD-95 expression per dendritic spine (which may result from upregulation of PSD-95 levels as well as elimination of small spines with low PSD-95 content). In a separate experiment we found that fear extinction-induced PSD-95 and dendritic spines changes were transient, as they were not observed 60 minutes after contextual fear extinction session, and they were specific for fear extinction, as we did not found them in the animals exposed to neutral context, as compared to the 5US group (Figure S2).

### The role of dCA1 PSD-95(S73) phosphorylation in regulation of fear extinction-induced PSD-95 expression

Based on the observed changes of PSD-95 levels and dendritic spines in dCA1 during contextual fear extinction, we hypothesized that extinction-induced upregulation of PSD-95 enables remodeling of the necessary circuits for contextual fear extinction memory. To validate this hypothesis, we used dCA1-targeted overexpression of phosphorylation-deficient PSD-95, with serine 73 mutated to alanine [PSD-95(S73A)]. We focused on serine 73 as its phosphorylation by αCaMKII negatively regulates activity-induced spine growth (Gardoni et al., 2006; Stein et al., 2003) and αCaMKII autophosphorylation-deficient mice have impaired contextual fear memory extinction (Radwanska et al., 2011). Accordingly, we expected that overexpression of PSD-95(S73A) would escalate fear extinction-induced accumulation of PSD-95 and spine growth. We did not use a phospho-mimetic form of PSD-95 (S73D), as this mutant protein locates mostly in dendrites in our hands (data not shown) and, therefore, unlikely affects synaptic function.

We designed and produced adeno-associated viral vectors, isotype 1 and 2 (AAV1/2) encoding mCherry under αCaMKII promoter (Control), wild-type PSD-95 protein fused with mCherry (AAV1/2:CaMKII_PSD-95(WT):mCherry) (WT) and phosphorylation-deficient PSD-95, where serine 73 was changed for alanine, fused with mCherry, (AAV1/2:CaMKII_PSD-95(S73A):mCherry) (S73A). We did not use a PSD-95 shRNA and shRNA-resistant PSD-95 genetic replacement strategy (Steiner et al., 2008) as these viruses depleted total PSD-95 levels *in vivo* in our hands (data not shown). The Control, WT and S73A viruses were stereotactically injected into the dCA1 of C57BL/6J mice (Figure 2A). Viral expression was limited to the dCA1 (Figure 2B). Expression of WT and S73A viruses resulted in significant overexpression of PSD-95 protein in three domains of a dendritic tree, compared to the Control virus (Figure 2C-D) (effect of virus: F(2, 30) = 13.09, P < 0.0001). Correlative light and electron microscopy confirmed that the overexpressed PSD-95 (WT and S73A) co-localised with postsynaptic densities (PSDs) of postsynaptic glutamatergic synapses; but weake signal was also present in dendrites (Figure 2E). Next, we investigated how fear extinction memory affects exogenous PSD-95 protein expression.

**Figure 2.**
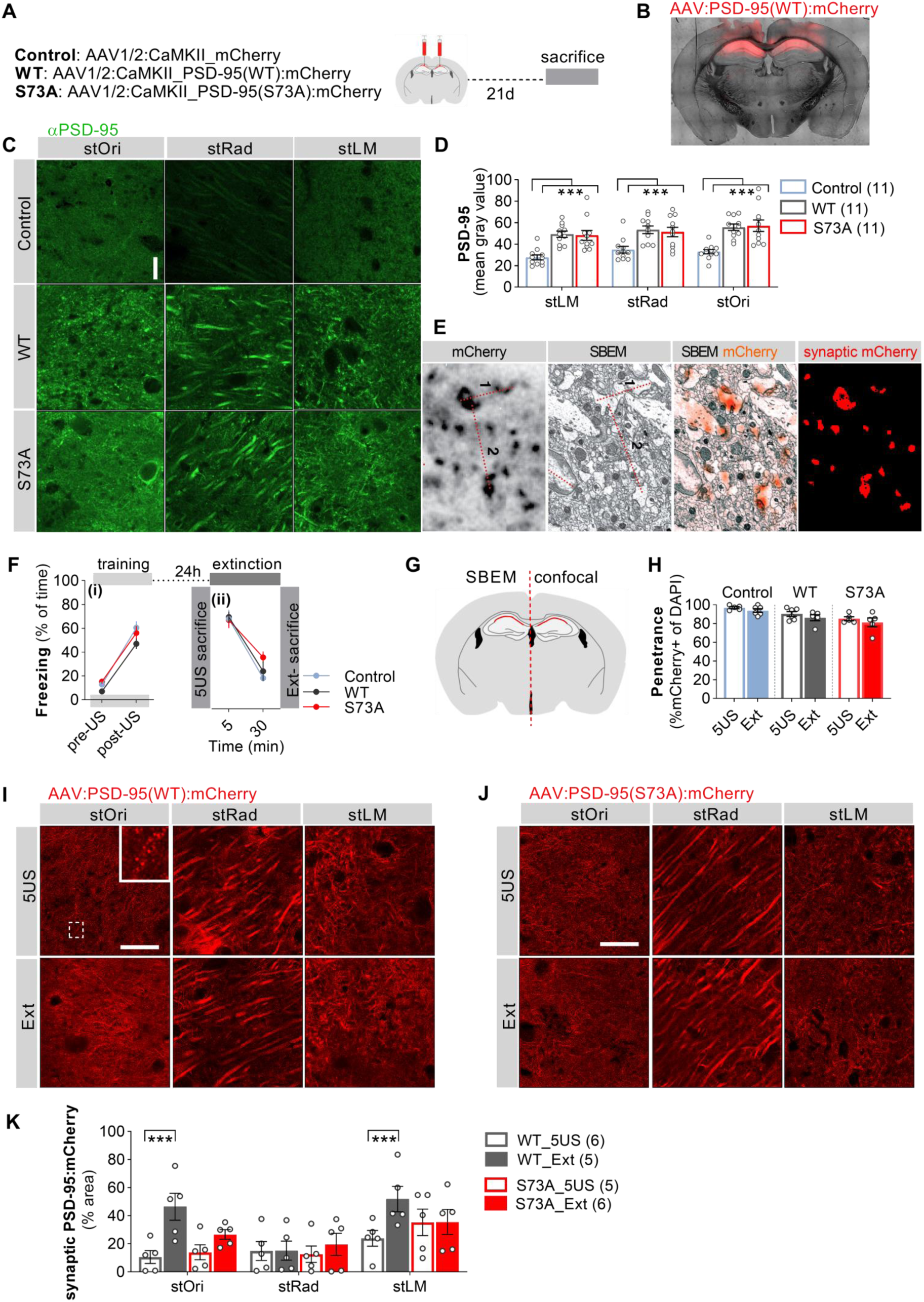
Phosphorylation of PSD-95(S73) is required for upregulation of PSD-95 levels during fear extinction training. **(A)** Mice were stereotactically injected in the dCA1 with AAV1/2 encoding Control (mCherry, n = 9), PSD-95(WT) (WT, ) or PSD-95(S73A) (S73A). **(B)** Microphotography of a brain with dCA1 PSD-95(WT):mCherry (WT) expression. **(C-D)** Analysis of PSD-95 overexpression. Representative confocal scans of the brain slices immunostained for PSD-95. Scale bar, 10 μm. Quantification of local expression of PSD-95 in three domains of dCA1 in mice with Control, WT and S73A. **(E)** Overexpressed WT co-localises with postsynaptic densities. Single confocal scan of overexpressed WT in dCA1, SBEM scan of the same area, superposition of confocal (orange) and SBEM images based on measured distances between large synapses (1 & 2), and thresholder synaptic WT signal. Measurements: (confocal image) 1: 3.12 μm, 2: 4.97 μm; (SBEM image) 1: 2.98 μm, 2: 4.97 μm. **(F)** Experimental timeline and percentage of freezing during **(i)** fear conditioning (5US) and **(ii)** fear extinction (Ext) session of mice with dCA1-targeted expression of Control, WT or S73A (mice: 5US/Ext, Control = 5/6; WT = 5/6; S73A = 5/5). **(G)** Illustration of the brain processing scheme**. (H)** Summary of data showing penetrance of the viruses in dCA1 (sections used for confocal and SBEM analysis). **(I-K)** Expression of the exogenous PSD-95 in dCA1. **(I-J)** Representative, confocal scans of the fused mCherry protein in three strata of dCA1. Inset: magnification of a dashed line rectangle. Scale bars, 10 μm. (**K**) Quantification of the PSD-95:mCherry-positive puncta (mice: 5US/Ext, Control = 5/6; WT = 5/6; S73A = 5/5). For C.ii, G and H.iii, each dot on the graphs represents one mouse (n indicated in the legends). ***P < 0.001.

A new cohort of mice with dCA1-targeted expression of the Control, WT and S73A underwent contextual fear conditioning (Figure 2F). Mice in all experimental groups showed increased freezing levels at the end of the training (RM ANOVA, effect of training: F(1, 30) = 269.4, P < 0.001, effect of virus: F(2, 30) = 2.815, P = 0.076) (Figure 2F). Half of the mice were sacrificed 24 hours after the fear conditioning (5US). The remaining half were re-exposed to the training box for fear extinction and sacrificed immediately afterward (Ext). All animals showed high freezing levels at the beginning of the session, which decreased during the session indicating extinction learning (RM ANOVA, effect of training: F(1, 15) = 65.68, P < 0.001). No effect of the virus was found (F(2, 15) = 0.993, P = 0.393) (Figure 2F).

For each animal, half of the brain was chosen at random for confocal analysis of the overexpressed PSD-95 protein, and the other half was processed for serial face-block scanning electron microscopy (SBEM) to analyse synapses at nanoscale resolution (Denk and Horstmann, 2004) (Figure 2G). The AAVs penetrance did not differ between the experimental groups (5US vs Ext) and reached over 80% in the analysed sections of dCA1 (Figure 2H). To assess the effect of the fear extinction session on the exogenous synaptic PSD-95 (WT and S73A) protein levels, we analysed fluorescent puncta formed by mCherry fused with PSD-95 protein that were small and intensive (Figure 2A, I and J). Three-way ANOVA indicated a significant effect of the training (F(1, 52) = 11.36, P = 0.0014) and dCA1 domain (F(2, 52) = 8.677, P = 0.006) on the expression of PSD-95:mCherry, but no effect of the virus (F (1, 52) = 0.8200, P = 0.369). *Post hoc* LSD analysis for the planned comparisons revealed that WT synaptic expression was upregulated in stOri (P = 0.016) and stLM (P = 0.035), but not stRad (P = 0.98), after the extinction session (Ext), compared to the 5US group (Figure 2K). Thus, the exogenous synaptic PSD-95(WT) protein levels were upregulated during fear extinction training in the same way as endogenous synaptic PSD-95. Surprisingly, no significant difference in the exogenous synaptic PSD-95(S73A) levels was observed between the Ext and 5US groups in all three strata of dCA1 (Figure 2K). Therefore, our data indicate that phosphorylation of PSD-95 at S73 is necessary for the fear extinction-induced upregulation of synaptic PSD-95 levels, although it does not affect the consolidation and recall of contextual fear memory or within-session reduction of fear.

### The role of PSD-95(S73) phosphorylation in regulating extinction-induced synapse remodeling

Since phosphorylation of PSD-95 at S73 is required for the fear extinction-induced upregulation of synaptic PSD-95, we hypothesized that PSD-95 also regulates extinction-induced synaptic growth. To test this, we used SBEM to determine dendritic spines density and to reconstruct spines and PSDs in the stOri (Figure 3A-C). PSDs are the postsynaptic elements that scale up with synaptic strength and are visible in electron microphotographs. In total, we reconstructed 159 spines from the brains of the mice expressing WT sacrificed 24 hours after contextual fear conditioning (5US) (n=3), and 178 spines from the mice sacrificed after fear extinction (Ext) (n=3). For mice expressing S73A, 183 spines were reconstructed in the 5US group (n=3) and 160 Ext (n=3). Lastly, we reconstructed 364 dendritic spines and PSDs in the Control 5US mice (n=3), and 293 spines from Ext (n =3). Figure 3D shows reconstructions of dendritic spines from representative SBEM brick scans for each experimental group.

**Figure 3.**
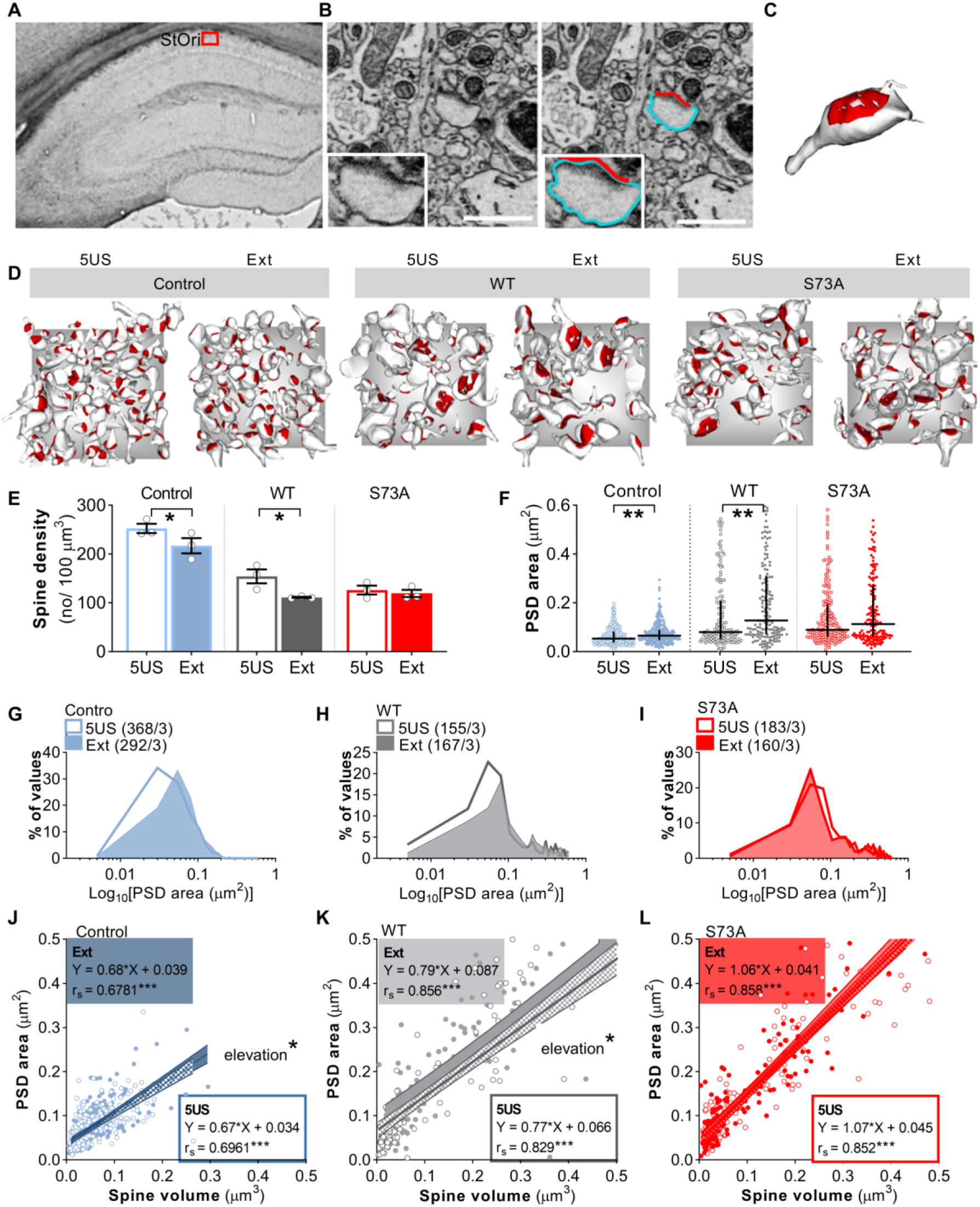
Phosphorylation of PSD-95 at S73 is required for synapse elimination and growth of remaining PSDs in stOri after fear extinction training. (**A-C**) The principles for SBEM analysis of the ultrastructure of dendritic spines and PSDs. **(A)** Microphotography of a dorsal hippocampus with the region of interest for analysis; **(B)** Tracing of a dendritic spine and PSD. Scale bars, 0.5 μm. A representative trace of a dendritic spine (in blue) and its PSD (in red), and **(C)** the reconstruction of this spine. **(D)** Exemplary reconstructions of dendritic spines and their PSDs from SBEM scans. The grey background rectangles are x = 4.3 × y = 4.184 μm. Dendritic spines and PSDs were reconstructed and analysed in tissue bricks. **(E)** Mean density of dendritic spines was downregulated after fear extinction (Ext) compared to trained (5US) Control and PSD-95(WT) (WT), but not PSD-95(S73A) (S73A) groups. **(F)** Median PSD surface area was increased after fear extinction (Ext) in Control and WT mice but did not change in S73A group. (**G-I**) Distributions were shifted towards bigger values in **(G)** Control and **(H)** WT groups **(I)** but not in the S73A group. X axes are Log_10_-transformed to show the differences between the groups. **(J-L)** Graphs showing changes in the correlation of dendritic spine volume and PSD surface area in **(J)** Control, **(K)** WT, **(L)** and S73A groups before (5US) and after extinction (Ext) training. For F, and J-L, each dot represents an individual dendritic spine; medians with IQR are shown. ***P < 0.001, **P < 0.01, *P < 0.05. Numbers of the analyzed dendritic spines/mice are indicated in (**G-I**).

Overexpression of PSD-95 protein (WT and S73A) resulted in decreased dendritic spines density and increased surface area of PSDs, compared to the Control group (Figure S3). We also observed a significant effect of the training on dendritic spines density (F(1, 45) = 8.01, P = 0.007). *Post hoc* analysis showed that the dendritic spines density was downregulated in the Control and WT Ext groups compared to their respective 5US groups (Fisher’s LSD test for planned comparisons, P < 0.035 and P < 0.014). No significant difference was observed for S73A Ext and 5US groups (Figure 3E). Furthermore, the median value of PSD surface areas was increased after the extinction training in the Control and WT groups (Mann-Whitney test, U = 42410, P < 0.001 and U = 9948, P < 0.001), but not in the S73A group (U = 13578, P = 0.246) (Figure 3F). The changes of PSDs surface area after extinction compared to 5US groups were also indicated as shifts in the frequency distribution toward bigger values in Control and WT groups (Figure 3G, H), but not in S73A (Figure 3I). We also observed the upward shift of the correlation lines of spine volume and PSD surface area after extinction training in Controls (ANCOVA, elevation: F(1, 6) = 4.677, P = 0.031) and WT groups (elevation: F(1, 319) = 4.256, P = 0.039), compared to their respective 5US groups (Figure 3J, K). Therefore, dendritic spines had relatively bigger PSDs after fear extinction than the dendritic spines of the same size in the 5US groups. Such a shift was not observed in the mice overexpressing S73A (elevation: F(1, 340) = 0.603, P = 0.437) (Figure 3L). Thus, in Control and WT groups, as in Thy1-GFP mice, elimination of dendritic spines observed after fear extinction was accompanied by an increased median area of the remaining synapses, indicating remodeling of the dCA1 circuits. The overexpression of S73A impaired both fear extinction-induced synaptic elimination and synaptic growth. We also confirmed the effect of PSD-95-overexpression and fear extinction training on synaptic transmission in dCA1 using *ex vivo* field recordings. We observed that after fear extinction the amplitude of field excitatory postsynaptic potentials (fEPSPs) was increased in the stOri dCA1 (when Shaffer collaterals were stimulated) of the mice that overexpressed PSD-95(WT), compared to their respective 5US groups (Figure S4), indicating enhanced excitatory synaptic transmission. Such change was not seen in S73A mice. There was also no effect of the extinction training on the fiber volley in both WT and S73A groups. Altogether, the electrophysiological analysis shows that PSD morphologic changes and functional alterations of synapses confirm the role of PSD-95 in remodeling of dCA1 circuits in contextual fear extinction.

### The role of dCA1 PSD-95(S73) phosphorylation in contextual fear extinction memory

Since overexpression of phosphorylation-deficient PSD-95(S73A) impaired extinction-induced expression of PSD-95 as well as structural and functional changes of synapses but did not affect within-session fear extinction, we hypothesised that PSD-95-dependent remodeling of synapses is necessary for consolidation of fear extinction memory. To test this hypothesis, two cohorts of mice with dCA1-targeted expression of the Control virus, WT or S73A, underwent contextual fear conditioning and fear extinction training. The first cohort underwent short extinction training with one 30-minut extinction session (Ext1) and 5-minut test of fear extinction memory (Test) (Figure 4D), while the second underwent extensive fear extinction training with three 30-minute contextual fear extinction sessions on the days 2, 3, 4 (Ext1-3), followed by spontaneous fear recovery/ remote fear memory test on day 18, and further four extinction sessions on the days 18, 19, 20, 21 (Ext4-7). Next, fear generalisation was tested in a context B (CtxB, day 22) (Figure 4E). The post-training analysis showed that the viruses were expressed in dCA1 (Figure 4A). The Control virus was expressed in 85% of the dCA1 cells, WT in 88% and S73A in 87% (Figure 4B-C). The analysis of short extinction training (data pooled from two cohorts) showed that in all experimental groups freezing levels were high at the beginning of Ext1 indicating a similar level of contextual fear memory acquisition (Figure 4D). However, freezing measured during the Test was significantly decreased, as compared to the beginning of Ext1, only in the Control (Fisher’s LSD for planned comparisons, P < 0.001) and WT (P = 0.004) groups, not in the S73A animals (P = 0.090) (RM ANOVA, effect of time: F(1, 46) = 26.13, P < 0.001, genotype: F(2, 46) = 0.540, P = 0.586; time x genotype: F(2, 46) = 1.25, P = 0.296). The analysis of freezing levels during the extensive fear extinction training also showed high levels of freezing at the beginning of Ext1 for all experimental groups (Figure 4E). In the Control and WT groups, the freezing levels decreased over consecutive extinction sessions (Ext2-6) and were significantly lower as compared to Ext1 (Fisher’s LSD for planned comparisons, P < 0.05 for all comparisons), indicating formation of long-term fear extinction memory (RM ANOVA, effect of time: F(3.681, 95.70) = 13.01, P < 0.001; genotype: F(2, 26) = 1.23, P = 0.306; time x genotype: F(10, 130) = 1.49, P = 0.147). We also found no spontaneous fear recovery after 14-day delay (Ext4 vs Ext3; Control, P = 0.806; WT, P = 0.248). In the S73A group, the extensive contextual fear extinction protocol did not reduce freezing levels measured at the beginning of Ext2-6 sessions, as compared to Ext1 (Fisher’s LSD for planned comparisons, P > 0.05 for all comparisons), indicating no fear extinction (Figure 4E). Accordingly we found significantly larger reduction of freezing during extensive fear extinction training (ΔExt6-Ext1) in the Controls (Tukey’s multiple comparisons test, P = 0.032) and WT animals (P = 0.026), as compared to the S73A group (one-way ANOVA, F(2, 24.94) = 4.98, P = 0.015) (Figure 4F). We also confirmed that the freezing reaction was specific for the training context, as it was very low and similar for all experimental groups in the context B (one-way ANOVA, F(2, 17.56) = 0.902, P = 0.424) (Figure 4G). Thus, our data indicate that overexpression of S73A in dCA1 does not affect fear memory formation, recall, or within-session extinction but prevents consolidation of contextual fear extinction memory even after extensive extinction training.

**Figure 4.**
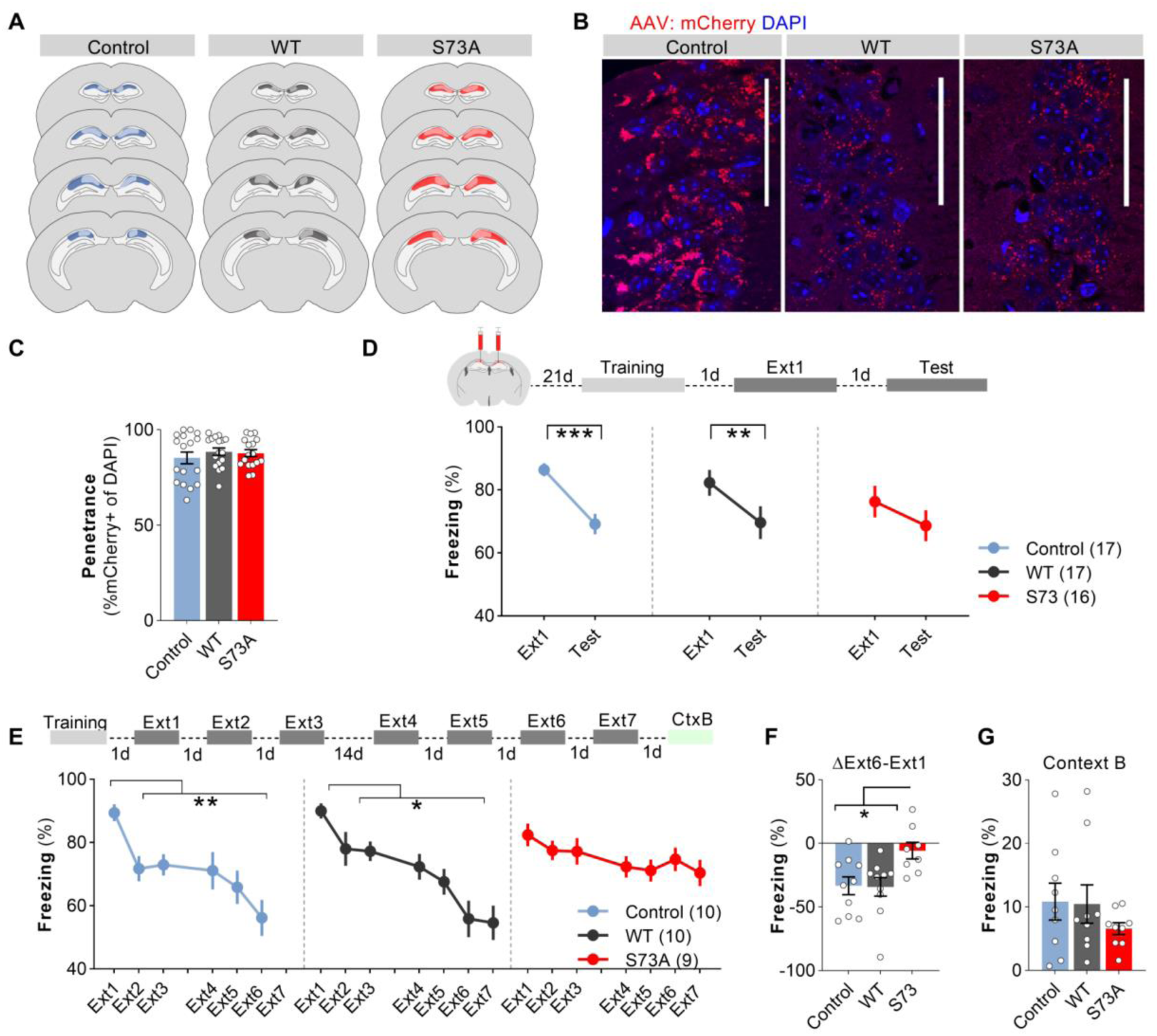
Phosphorylation of PSD-95 at serine 73 is required for contextual fear extinction. **A,** Area and extent of viral infection shown. **B,** Single confocal scans of the stratum pyramidale of dCA1 of the mice expressing Control (n = 17), WT (n = 17) and S73A (n = 16) (scale bars, 50 μm) and **(C)** penetrance of the viruses. **D.** Experimental timeline and percentage of freezing during fear extinction and consolidation of fear extinction memory test of the mice with dCA1-targeted expression of Control, WT or S73A. **E-G.** Experimental timeline and percentage of freezing during extensive fear extinction training of the mice with dCA1-targeted expression of Control (n=10), WT (n=10) or S73A (n=9). **F.** Summary of data showing change of freezing levels during extensive fear extinction training, as compared to the Ext1, and (**G)** the test of fear levels in the context B. *P < 0.05; **P < 0.01; ***P < 0.001 by Tukey’s multiple comparisons tests.

### Effect of chemogenetic inhibition of dCA1 on fear extinction-induced expression of PSD-95

Our data indicate that PSD-95(S73A) overexpression prevents extinction-induced upregulation of PSD-95 and synaptic remodeling, as well as the extinction of fear memory. These observations suggest that extinction-induced upregulation of PSD-95 is required to update an extinguished fear memory. However, behavioral impairments induced by overexpression of S73A may result from the deregulation of PSD-95 levels at other time points of training. Accordingly, we asked whether the dCA1 activity, specifically during the first extinction session, is required for extinction-induced PSD-95 expression. Such findings would support the hypothesis that extinction-induced PSD-95 expression is required to extinguish fear memory.

To test this hypothesis we used chemogenetic tools to manipulate dCA1 activation during the fear extinction memory session and analysed extinction-induced PSD-95 expression. AAV1/2 encoding inhibitory designer receptors exclusively activated by designer drugs (DREADD, hM4(Gi)) under human synapsin (hSyn) promoter [AAV1:hSyn-hM4(Gi):mCherry (hM4)] (Roth, 2016), or a Control virus encoding mCherry (AAV1/2:CaMKII-mCherry) were bilaterally infused into the dCA1 region of mice. The post-training analysis of the hippocampal sections confirmed that the expression of the viruses was limited to the dCA1 (Bregma > -2.5 mm) (Figure 5A). There were no significant differences in the virus penetration between the experimental groups [hM4 was expressed in 71% and 80% of the pyramidal cells (in the saline and CNO groups, respectively); the Control virus was expressed in 84% and 87% of the cells (saline and CNO, respectively)] (Figure 5B). Both groups of the mice with hM4 virus showed low freezing levels at the beginning of the training session, and freezing increased after USs delivery (RM ANOVA, effect of time: F(1, 10) = 86.36, P < 0.0001) (Figure 5C). The next day, mice received a systemic injection of saline or CNO (1 mg/kg), and 30 minutes later, they were re-exposed to the training context. As in previous experiments, both groups of mice showed high levels of freezing at the beginning of the extinction session, which decreased throughout the session (effect of time: F(1, 11) = 8.149, P = 0.016), indicating within-session extinction. There was no effect of drug (F(3, 26) = 2.438, P = 0.087), or a training and drug interaction (F(3, 26) = 1.086; P = 0.372), on the freezing levels (Figure 5C). At the end of the 30-minute extinction session, the brains were collected and immunostained to detect PSD-95 protein (Figure 5D). There was a significant effect of the drug (F(1, 16) = 31.06, P < 0.0001), but no effect of the CA1 domain (F(2, 29) = 0.739, P = 0.486), on PSD-95 levels. *Post hoc* LSD tests for planned comparisons confirmed that the expression of PSD-95 was decreased in all strata of dCA1 in the CNO group, compared to the saline-treated animals (P < 0.05 for all domains) (Figure 5E). To validate whether this downregulation of PSD-95 expression was specific to the chemogenetic inhibition, we trained mice with the Control virus expressed in the dCA1 (Figure 5F). The animals were injected with CNO before the extinction session and sacrificed after the session (Figure 5F). As in the previous experiment, CNO did not affect memory recall or within-session fear extinction (effect of drug: F(3, 27) = 1.628, P = 0.206). Moreover, there was no significant effect of the drug (RM ANOVA, effect of drug: F(1, 12) = 3.73, P = 0.077) or the region (F(1.302, 14.32) = 1.505, P = 0.248) on PSD-95 expression levels (Figure 5G-H), indicating that CNO does not affect PSD-95 expression.

**Figure 5.**
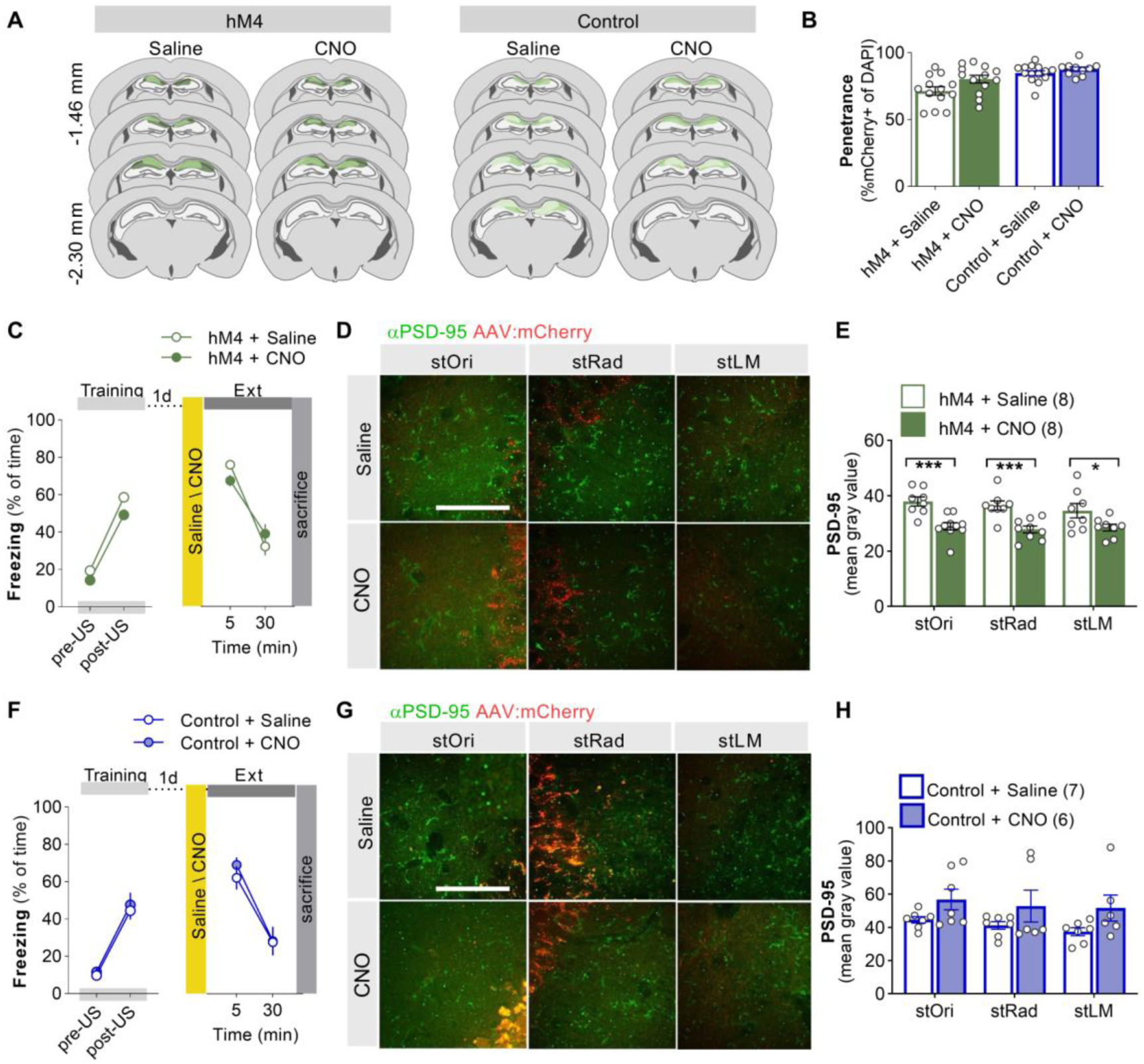
Chemogenetic inhibition of dCA1 during extinction session impairs extinction-induced PSD-95 expression. Three weeks after bilateral dCA1 viral infusion surgery, mice were trained in fear conditioning, followed by a fear extinction session 24 hours later. In all groups, saline or CNO was systemically injected 30 minutes before the extinction session. Mice were sacrificed immediately after the fear extinction session. **(A)** The extent of viral transfections in the CA1 area. **(B)** Penetrance of hM4 virus and Control virus in dCA1(mice, saline/CNO, hM4 = 8/8; Control = 7/6). **(C)** Experimental timeline and percentage of freezing during fear conditioning and fear extinction session of the mice with hM4 virus. **(D)** Representative, confocal scans of the brain slices immunostained for PSD-95 in hM4 groups. Scale bar: 10 μm. **(E)** Summary of data quantifying the expression of PSD-95 in three domains of dCA1 of the mice with hM4 virus. **(F)** Experimental timeline and percentage of freezing during fear conditioning and fear extinction session of the mice with the Control virus. **(G)** Representative, confocal scans of the brain slices immunostained for PSD-95. Scale bar: 10 μm. **(H)** Summary of data quantifying the expression of PSD-95 in three domains of dCA1 of the mice with the Control virus. *P < 0.05, **P < 0.01; ***P < 0.001.

Since dendritic spines in dCA1 undergo constant remodeling (Attardo et al., 2015) the effect of the chemogenetic inhibition of dCA1 neurons on PSD-95 levels could be unrelated to the extinction-induced PSD-95 expression but results from decreased cell activity. To test this hypothesis we chemogeneticaly inhibited dCA1 neurons outside of the fear extinction time-window (7d after training) and measured the changes of PSD-95. Mice with bilateral expression of hM4 or the Control virus were systemically injected with saline or CNO (Figure S5). As in the extinction experiment, the brains were collected 60 minutes after the injection and immunostained for PSD-95. At this time point, no effect of the drug on PSD-95 levels was observed in the Control or hM4 groups (Figure S5). Thus, chemogenetic inhibition of dCA1 outside of the fear extinction memory window does not affect the levels of PSD-95.

### The effect of chemogenetic inhibition of dCA1 area during fear extinction on updating an extinguished contextual fear memory

Our experiments showed that chemogenetic inhibition of dCA1 during extinction of contextual fear memory prevented the extinction-induced expression of PSD-95. Thus extinction-induced upregulation of PSD-95 levels in the dCA1 is a likely mechanism that enables extinction of contextual fear memory. To test this hypothesis, we again used chemogenetic tools. Mice were bilaterally injected in the dCA1 with AAV1/2 encoding hM4 or the Control virus, and they were trained 3 weeks later (Figure 6A). The post-training analysis of the hippocampal sections revealed that hM4 was expressed in 76% of the pyramidal cells of dCA1 (both in cell bodies and dendrites), while the Control virus in 84% of the cells (Figure 6B, C). The expression of the virus was limited to the dCA1 (Bregma > -2.5 mm) (Figure 6D). Three weeks post-surgery and viral infection, mice underwent contextual fear conditioning. Twenty-four hours after training, mice received a systemic injection of saline or CNO (1 mg/kg) to activate hM4 receptors, and were re-exposed to the training context for contextual fear extinction (Ext) (Figure 6E). Mice in all groups showed high freezing levels at the beginning of Ext, indicating fear memory formation and no drug-induced impairment of memory recall (Figure 6E). We next tested fear extinction memory 24 hours later (Test). Only in the hM4 group injected with saline, but not in the group injected with CNO, the freezing levels during the Test were lower as compared to Ext (RM ANOVA, effect of time: F(1, 32) = 11.22, P = 0.002, drug: F(1, 32) = 0.112, P = 0.739; LSD *post hoc* tests for planned comparisons, Saline: P < 0.001; CNO: P = 0.214), indicating consolidation of fear extinction memory in the hM4+Saline group and impairment of fear extinction by chemogenetic inhibition of dCA1 (Figure 6E). In the Control virus groups, the freezing levels decreased during the Test as compared to Ext, and no effect of the drug was observed (effect of time: F(1, 24) = 24.2, P < 0.001; drug: F(1, 24) = 1.29, P = 0.267; LSD *post hoc* tests for planned comparisons, Saline: P < 0.001; CNO: P = 0.005) (Figure 6F). Thus, CNO alone did not impair consolidation of fear extinction memory. Therefore, we next asked whether chemogenetic manipulation of dCA1 outside (a day prior) the fear extinction memory session (Ext) impairs updating of the fear memory.

**Figure 6.**
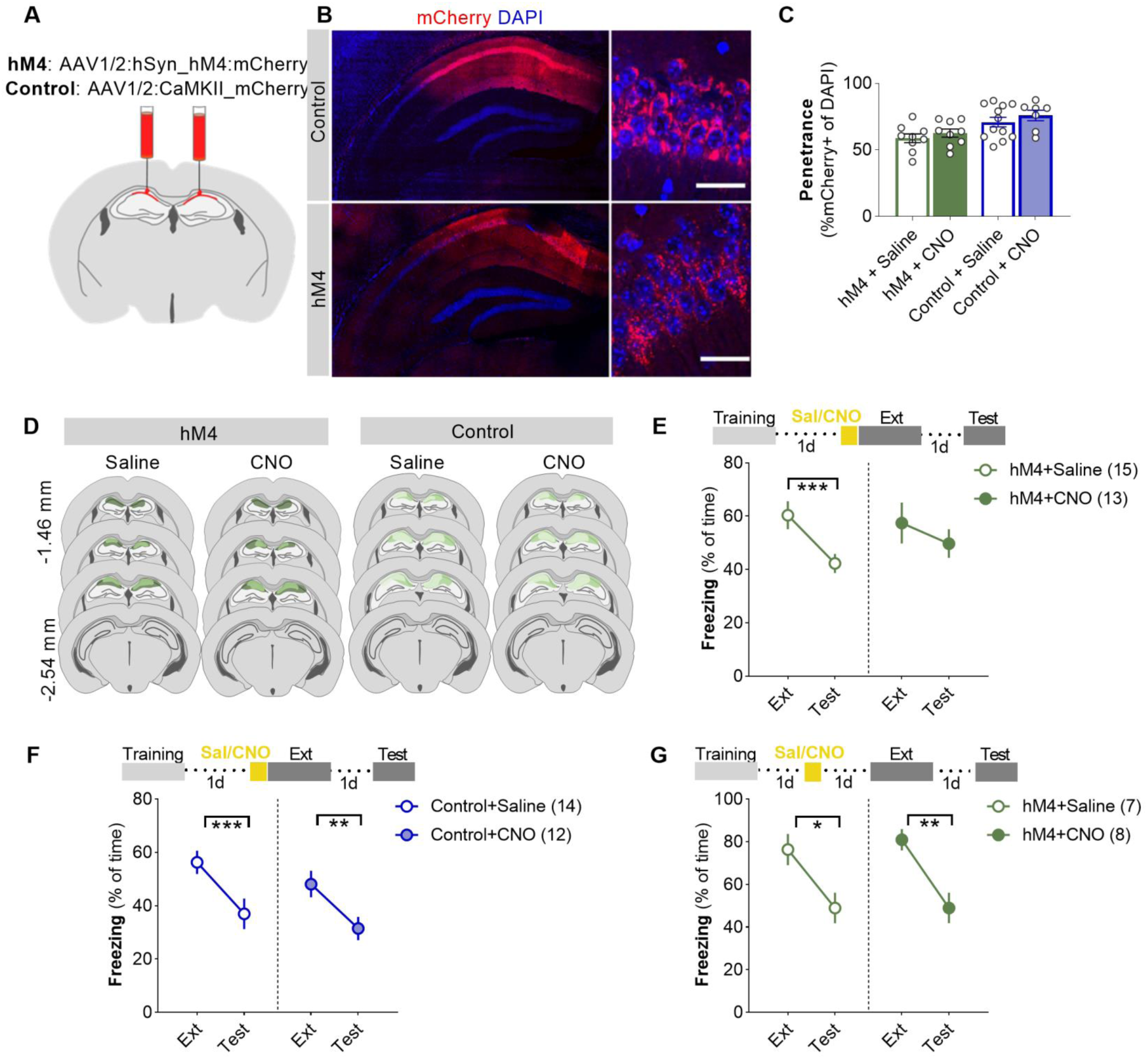
Chemogenetic inhibition of dCA1 impairs extinction of contextual fear. **(A)** AAV-hSyn-hM4(Gi):mCherry (hM4) and Control virus (mCherry) were bilaterally injected into the dCA1 (left). **(B)** Single confocal scans showing transfected dCA1. Viral expression was observed in cell bodies and dendrites of pyramidal neurons. (right) Magnification of dCA1 confocal scans (Scale bars, 30 μm). (**C**) Penetrance of hM4 virus (n = 27 animals) and Control virus (n = 24 animals). **(D)** The extent of viral transfections in the CA1 area. Minimum and maximum transfections are shown. (**E-F**) Experimental design and percentage of freezing during fear extinction session (Ext) and consolidation of fear extinction memory test (Test) of the mice with hM4 **(E)** and Control virus **(F)**. Mice were trained three weeks after the surgery for optimal virus expression. In all groups, saline or CNO (1mg/kg) was systemically injected 30 minutes before Ext. **(G)** Experimental design and percentage of freezing during fear extinction session (Ext) and Test of the mice with hM4 virus. Mice were trained three weeks after the surgery and virus expression. CNO or Saline was systemically injected 24 hours before extinction (Ext). ***P < 0.001, **P < 0.01, *P < 0.05 by LSD test for planned comparisons only, Ext vs. Test. Numbers of trained mice are indicated in the legends.

A new group of C57BL/6J mice were injected into dCA1 with AAV1/2 encoding hM4 and trained 3 weeks later (Figure 6G). The virus penetrance and area of the infection were similar to previous experiments. One day after training, all mice received a systemic injection of CNO (1 mg/kg) or Saline and were re-exposed to the training context 24 hours later for fear extinction (Ext). On the following day fear extinction memory was tested (Test). Mice from both groups showed high freezing levels at the beginning of Ext, and it was lower during Test as compared to Ext (RM ANOVA, effect of training: F(1, 14) = 270, P < 0.0001; drug: F(1, 15) = 0.134, P = 0.719; LSD *post hoc* tests for planned comparisons, Saline: P = 0.014; CNO: P = 0.003), indicating no impairment of fear memory recall and extinction (Figure 6G). Overall, our data indicate that chemogenetic inhibition of dCA1 during the contextual fear extinction session does not affect fear memory recall but prevents extinction of the contextual fear memory leading to fear memory persistence.

## DISCUSSION

Here, we have investigated synaptic processes in the dCA1 that contribute to contextual fear memory attenuation. Our interest in this problem stems from many anxiety disorders associated with impaired fear extinction and hippocampus function (van Rooij et al., 2018). The key findings from the present study are that (1) contextual fear extinction increases PSD-95 protein expression per dendritic spine in the dCA1 and is accompanied by remodeling of the glutamatergic synapses; (2) this extinction-induced PSD-95 expression and synaptic remodeling is regulated by phosphorylation of PSD-95 at serine 73; (3) PSD-95 phosphorylation at serine 73 in the dCA1 is required for extinction of fear memories but not for the fear memory consolidation or recall. Below, the significance of the findings is discussed in light of previous studies.

PSD-95 affects the structure and function of glutamatergic synapses. In particular, *in vitro* studies showed PSD-95 overexpression increases the size of glutamatergic synapses (Nikonenko et al., 2008). Our study is the first to show how overexpression of PSD-95 influences dCA1 glutamatergic synapses *in vivo*. We confirm that the overexpression of PSD-95 (both WT and S73A) increases the median areas of PSDs, and it also results in a loss of small dendritic spines. Thus, the structural consequences of PSD-95 overexpression *in vivo* are profound as they involve the global remodeling of the local circuit, but the long-term elimination and up-scaling of synapses are not regulated by PSD-95 serine 73 phosphorylation as similar changes are observed in WT and S73A 5US groups, as compared to the Control 5US. Moreover, we demonstrate that contextual fear extinction induces rapid loss of synapses in the stOri that is accompanied by heterosynaptic upregulation of PSD-95 levels, growth of the synapses and increased synaptic transmission. Upregulation of PSD-95 levels during memory formation and recall was previously demonstrated in the hippocampus and cortex (Elkobi et al., 2008; Zanca et al., 2019). Here, it is likely that protein translation, degradation, translocation as well as the loss of small spines with low PSD-95 content contribute to the relative upregulation of PSD-95 levels per dendritic spine in the Ext group, as compared to 5US animals.

The synaptic processes induced by fear extinction allude to the Hebbian strengthening of activated synapses and heterosynaptic weakening of adjacent synapses observed in activated visual cortex neurons and *in vitro* (El-Boustani et al., 2018; Royer and Paré, 2003). Our study is the first description of bidirectional plasticity of dendritic spines in the dCA1 during attenuation of fear memories. Previously, the heterosynaptic weakening was shown to be driven by the expression of CaMKII-regulated Arc protein (El-Boustani et al., 2018). Here, we show that both aspects of the fear extinction-induced synaptic plasticity (spine elimination and growth of the remaining synapses) are coordinated by αCaMKII-dependent phosphorylation of PSD-95 at serine 73 (Gardoni et al., 2006). This is a new function of PSD-95 serine 73 as previously it was shown to be required for: PSD-95 dissociation from the NMDAR subunit NR2A after NMDAR stimulation (Gardoni et al., 2006), PSD-95 protein downregulation during LTD (Nowacka et al., 2020) and termination of synaptic growth after glutamate uncaging (Steiner et al., 2008). Thus none of these synaptic models explains synaptic processes observed during fear extinction as they predict excessive growth of the synapses and accumulation of PSD-95(S73A). The effects of S73A mutation can be explained assuming interdependence of bidirectional synaptic processes induced by fear extinction; for example, synaptic growth is only allowed if some synapses are eliminated (e.g. due to spatial constraints), and the later process is precluded due to stable PSD-95(S73A)-NMDAR interactions at PSD (Gardoni et al., 2006). The precise timing and location of dCA1 PSD-95(S73) phosphorylation and dissociation of PSD-95-NMDAR complex to enable PSD elimination during fear extinction has to be revealed in the future studies.

Our data indicate that the extinction of contextual fear induces the upregulation of PSD-95 expression per dendritic in the stOri and stLM, while the protein levels in stRad are not changed. These alterations are accompanied by the increased median area of PSDs, indicating circuit remodeling in the distal strata of dendrites. As shown by the control experiments, the extinction-induced synaptic changes are transient (not observed 60 minutes after contextual fear extinction session), and absent in the animals exposed to neutral and known context (without USs experience during training) proving their specificity for fear extinction. Interestingly, chemogenetic inhibition of dCA1 during fear extinction session downregulates PSD-95 also in stRad suggesting that, although the net changes of PSD-95 levels are not detected, these synapses also are remodelled but to lesser extent. Thus the extinction-induced synaptic change pattern is strikingly different from the changes observed immediately after contextual fear memory encoding where transient synaptogenesis is observed in the stRad (Radwanska et al., 2011). These observations support the idea that different CA1 inputs are involved in memory formation and extinction. CA3 neurons project to the stRad and stOri regions of CA1 pyramidal neurons, the nucleus reuniens (Re) projects to the stOri and stLM, and the entorhinal cortex (EC) projects to the stLM (Hoover and Vertes, 2012; Ishizuka et al., 1990; Kajiwara et al., 2008; Vertes et al., 2015). Thus, the pattern of synaptic changes induced by contextual fear extinction co-localises with the domains innervated by the Re and EC, suggesting that these inputs are regulated during contextual fear extinction. In agreement with our observations, previous data showed that the EC is activated during and required for contextual fear extinction in animal models (Baldi and Bucherelli, 2015, 2014; Bevilaqua et al., 2006). Human studies also showed that EC-CA1 projections are activated by cognitive prediction error (that may drive memory extinction), while CA3-CA1 projections are activated by memory recall without prediction errors (Bein et al., 2020). The role of the Re in fear memory encoding, retrieval, extinction and generalisation has been demonstrated (Ramanathan et al., 2018; Troyner and Bertoglio, 2021; Xu and Sudhof, 2013). Still, it has to be established whether the plasticity of dCA1 synapses is specific to Re and/or EC projections.

The formation of spatial and contextual fear memories is thought to involve NMDA receptor-dependent synaptic plasticity in the dCA1 (Bliss and Collingridge, 1993; Lisman et al., 2017; Martin et al., 2000). However, more recent dCA1-targeted genetic manipulation studies have shown that mice with dCA1 knockout of NMDA receptor (NMDAR) subunit, NR1, have an intact formation of spatial and contextual fear memories (Bannerman et al., 2012; Hirsch et al., 2015). However, NMDAR-dependent synaptic transmission is required for spatial choice (Bannerman et al., 2012) and contextual fear extinction (Hirsch et al., 2015). Accordingly, it has been proposed that NMDAR-dependent plasticity in the dCA1 has a crucial role in detecting and resolving contradictory or ambiguous memories when spatial information is required (Bannerman et al., 2014). For example, dCA1 NMDAR-dependent plasticity would be required during extinction training of contextual fear memories, in which an animal recalls aversive memories of the context (or cues) and experiences a conflicting new experience of the same context being safe. This is consistent with comparator views of hippocampal function (Gray, 1982; Grossberg and Merrill, 1992) and the fact that hippocampus processes surprising events such as novelty and prediction errors (Bein et al., 2020; Huh et al., 2009; Kumaran and Maguire, 2006; Ploghaus et al., 2000). In agreement with this theory, our experiments are the first to show that dCA1-targeted genetic manipulation blocking the phosphorylation of PSD-95 at serine 73, and chemogenetic inhibition during the fear extinction session, prevents fear extinction-induced dCA1 synaptic remodeling and extinction of contextual fear even after extensive extinction training. dCA1 PSD-95(S73A) mutation impairs extinction not only of recent (1-day old) but also remote (14-day old) contextual fear memory. We also show that dCA1 PSD-95(S73A) mutation does not affect mice activity, context-independent fear generalisation or fear recovery after 14-day delay. Thus our data support the hypothesis that PSD-95(S73)-dependent synaptic plasticity of the dCA1 is necessary to resolve conflicting pieces of information about the fear-associated context, and this refers to contextual information independent of its age and extent of novel and conflicting experience exposure. In agreement with our findings, Cai with collaborators (2018) and Li with collaborators (2017) show that the signaling pathways downstream of NMDAR-PSD-95 complex in the dorsal CA3 and DG are involved in contextual fear extinction. In particular, translocation of PSD-95 from NMDAR to TrkB, and increased PSD-95-TrkB interactions, promotes extinction, while competing NMDAR-PSD-95-nNOS interactions hinder contextual fear extinction by inhibiting ERK signalling (Cai et al., 2018) that is required for fear extinction (Tronson et al., 2009). Accordingly, PSD-95(S73A) mutation, that hampers dissociation of PSD-95 from NMDAR (Gardoni et al., 2006), may limit interactions of PSD-95 with TrkB, and therefore obstruct fear extinction. This adds up to previous studies investigating the molecular processes in dCA1, including activation of ERK, CB1, and CBEP, that are required for contextual fear extinction, but not fear memory consolidation (Berger-Sweeney et al., 2006; Bitencourt et al., 2008; de Oliveira Alvares et al., 2008; Pamplona et al., 2008; Radulovic and Tronson, 2010; Tronson et al., 2009). Interestingly, other processes, such as protein synthesis and c-Fos expression, are necessary for contextual fear consolidation and reconsolidation, but not extinction (Fischer, 2004; Lattal and Abel, 2004; Mamiya et al., 2009; Tronson et al., 2009). Thus, it remains puzzling how synaptic plasticity, without concomitant translation, contributes to contextual fear extinction.

In our study the local genetic and chemogenetic manipulations tend to decrease contextual fear memory retrieval (Ext1). However, the differences between the experimental groups never reach statistical significance. This observation is in agreement with a previous report (Hirsch et al., 2015), but contradicts other studies which found that genetic, optogenetic or excitotoxic inactivation of dCA prevents recall of contextual fear memory (Ji and Maren, 2008; Nagura et al., 2012; Sakaguchi et al., 2015). The methodological differences between ours and previous studies may explain discrepant results. Sakaguchi et al. (2015) used optogenetic stimulation of dCA1 in α-CaMKII-tTA × TetO-ArchT-GFP mice that expressed ArchT not only in the dCA1 neurons but also neurons that project to dCA1. Firstly, optogenetic inhibition affects not only synaptic transmission but also cell excitability. Secondly, we used more intensive behavioural training (5US vs 1 US), that results in memory which is more resistant to disruption (Irvine et al., 2005; Radwanska et al., 2011). Ji and Maren (2008) used excitotoxic inactivation of dCA1 and investigated cued fear conditioning and fear extinction to the cue. Thus context is only the background in their study and it may differently involve dCA1 synaptic plasticity than context used as a foreground factor. Furthermore, excitotoxic lesion, as optogenetic inhibition, affects not only synapses but also cell activity. Finally, Nagura and colleagues (2012) used ligand binding-deficient PSD-95 cDNA knockin (KI) mice and observed enhanced contextual fear memory formation and impaired long-term memory retention as in the following study (Fitzgerald et al., 2015). However, even though the behavioral phenotype was supported by ephys data showing impaired LTP in dCA1, it is unknown whether it really relies on CA1 plasticity. Thus, the analysed literature and our results support the notion that dCA1 synaptic plasticity is involved in contextual fear memory extinction, but it is not necessary for contextual fear memory retrieval. In particular, phosphorylation of PSD-95(S73) is not critical for fear memory formation and expression. As earlier EM studies show, contextual fear memory formation involves transient (< 24 hr) synaptogenesis in dCA1 (Radwanska et al., 2011), while here we demonstrate that contextual fear memory extinction involves elimination of dendritic spines and parallel growth of the remaining synapses (both of these phenomena being impaired by S73A mutation). Accordingly, we can propose that memory formation and simple synaptic strengthening (that is not coupled with dendritic spine elimination) are independent of PSD-95(S73), as previously shown (Steiner et al., 2008). However PSD-95(S73) is required for fear extinction and bidirectional plasticity induced in dCA1 during fear extinction.

### Conclusions

Our study pinpoints a cellular mechanism that operates in the dCA1 area and contributes to contextual fear memory attenuation. We propose that the propensity for extinction of contextual fear memories relies on opposing synaptic processes: strengthening of synapses and rapid elimination of small dendritic spines, that both require PSD-95 serine 73 phosphorylation. Since new or long-lasting memories may be repeatedly reorganized upon recall (Nader et al., 2000; Schafe et al., 2001), the molecular and cellular mechanisms involved in extinction of the existing fearful memories provide excellent targets for fear memory impairment therapies. In particular, understanding the mechanisms that underlie contextual fear extinction may be relevant for post-traumatic stress disorder treatment.

## Acknowledgments, Funding and Disclosure

This work was supported by a National Science Centre (Poland) Grant No. 2015/19/B/NZ4/02996 and 2013/08/W/NZ4/00861 to KR. PRELUDIUM Grant No. 2016/21/N/NZ4/03304 to MZ and PRELUDIUM Grant No. 2015/19/N/NZ4/03611 to KŁ. TW was supported by the National Science Centre (Poland) (Grant No. 2017/26/E/NZ4/00637). The project was carried out using CePT infrastructure financed by the European Union - The European Regional Development Fund within the Operational Program “Innovative economy” for 2007-2013.

MZ, MB, KFT and KR designed the experiments; MZ, MB, MNS, MR, AN, AC, KTF, AS, KŁ, TW and MŚ performed the experiments; MZ, MB, ES, MŚ, MR, KŁ, KFT, TB, JW and KR analyzed data. MZ, MB and KR drafted the manuscript. All authors had critical input to the final version of the manuscript. Authors report no financial interests or conflicts of interest. Light and microscopy experiments were performed at the Laboratory of Imaging Tissue Structure and Functions.

## MATERIALS AND METHODS

A full description of Materials and Methods are available in supplementary material online.

### Animals

C57BL/6J and Thy1-GFP(M) (Feng et al., 2009b) mice were used in the experiments. The experiments were undertaken in accordance with the Animal Protection Act of Poland and approved by the I Local Ethics Committee (261/2012, Warsaw, Poland).

### Contextual fear conditioning

The animals were trained in a conditioning chamber (Med Associates Inc, St Albans, USA) in a soundproof box. Mice were placed in the training chamber, and after a 148 s introductory period, a foot shock (2 s, 0.7 mA) was presented. The shock was repeated 5 times at 90 s inter-trial intervals. Contextual fear memory was tested and extinguished 24 h after training by re-exposing mice to the conditioning chamber for 30 minutes without US presentation, followed by a second 30-minute extinction session the following day. Freezing and locomotor activity of mice was automatically scored. In all experiments, experimenters were blind to the experimental groups.

## Supplementary Materials

## SUPPLEMENTARY RESULTS

**Supplementary Figure 1.**
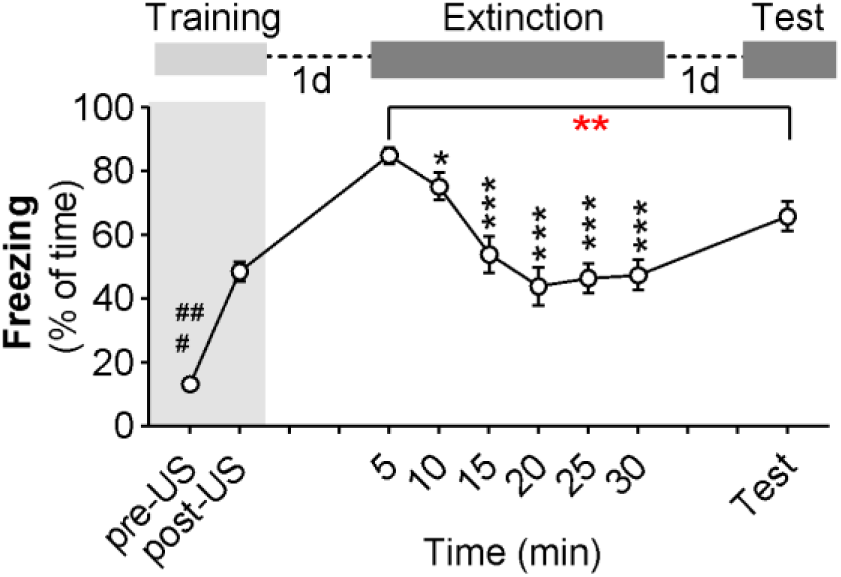
Experimental timeline and the freezing levels of the mice during contextual fear conditioning training, fear extinction session and fear extinction memory test. C57BL/6j mice (n = 15) showed low freezing levels in a novel context before delivery of electric shocks (pre-US) and freezing increased during the training (post-US) (t = 14.91, df = 14, **^###^**P < 0.001), indicating fear memory formation. Twenty-four hours later, the animals were re-exposed to the training context without US presentation for the fear extinction memory session (Extinction). Freezing levels were high at the beginning of the session, indicating fear memory retrieval and decreased within the session (RM ANOVA with Holm-Sidak’s multiple comparisons tests (black asterisks), F(3.011, 42.15) = 20.72, P < 0.001) indicating the formation of fear memory extinction. Next, we tested the consolidation of fear extinction memory 24 hours later (Test). At the beginning of Test, the freezing levels were lower than at the beginning of extinction 1 (t = 3.843, df = 14, **\*\***P = 0.0018), indicating the formation of long-term fear extinction memory.

**Supplementary Figure 2.**
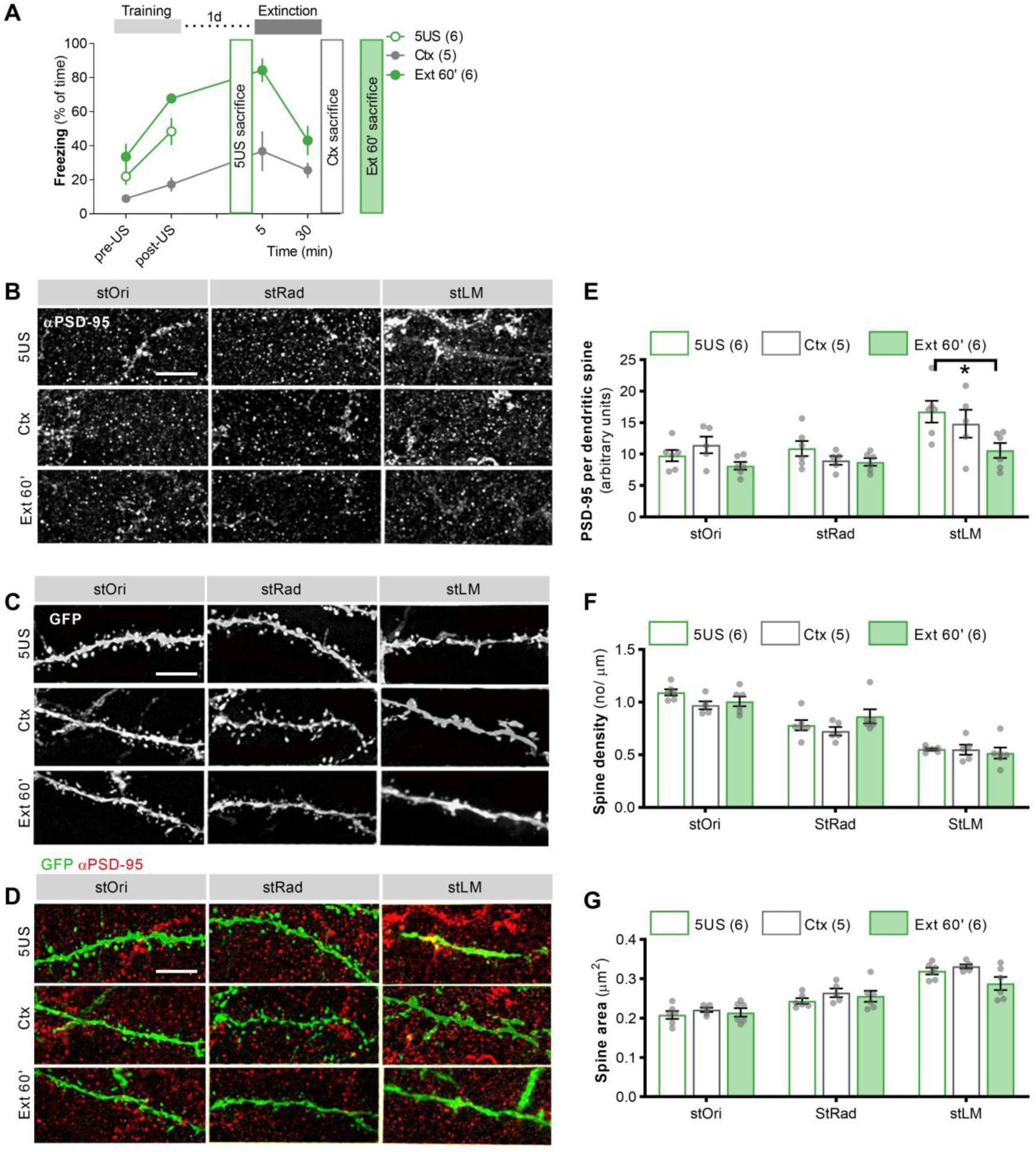
Fear extinction-induced PSD-95 and dendritic spines changes were transient and could not be induced by the exposure to neutral context. Dendritic spines were analysed in three domains of dendritic tree of dCA1 area in Thy1-GFP(M) mice: stOri, stRad and stLM. **(A)** Experimental timeline and freezing levels of mice from three experimental groups: fear conditioning training (5US, mice sacrificed 1 day after contextual fear conditioning; n = 6), context (Ctx, mice sacrificed immediatelly after the second exposure to novel context, no USs were delivered) and fear extinction 60’ (Ext 60’, mice sacrificed 60 minutes after contextual fear extincion session, n=6). **(B-D)** Representative confocal images of PSD-95 immunostaining. Thy1-GFP and their colocalization (maximum projections of z-stacks composed of 20 scans) are shown for three domains of the dendritic tree. (**E**) Summary of data showing PSD-95 expression per dendritic spine in stOri, stRad and stLM (mouse: 5US = 6; Ctx = 5; Ext 60’ = 6). There was no effect of training (F(2, 14) = 2.799, P = 0.095), but a significant effect of dendritic domain (F(1.574, 22.04) = 32.00, P < 0.001) and training x dendritic domain interaction (F(4, 28) = 4.191, P = 0.009). *Post hoc* Tukey’s test showed that PSD-95 area per dendritic spine was decreased in stLM in Ext 60’ group as compared to the 5US animals (P = 0.039). **(F)** Summary of data showing dendritic spines density. There was no effect of training (F(2, 14) = 1.620, P = 0.233), but a significant effect of dendritic domain (F(1.874, 26.23) = 79.64, P < 0.001), and no training x dendritic domain interaction (F(4, 28) = 1.43, P = 0.250). **(G)** Summary of data showing average dendritic spine area. There was no effect of training (F(2, 14) = 3.162, P = 0.074), but a significant effect of dendritic domain (F(1.340, 18.76) = 56.36, P < 0.001), and no training x dendritic domain interaction (F(4, 28) = 1.33, P = 0.283). For E-G each dot represents one mouse. Scale bars: E, G, H: 15 μm. *P < 0.05, **P < 0.01; ***P < 0.001.

**Supplementary Figure 3.**
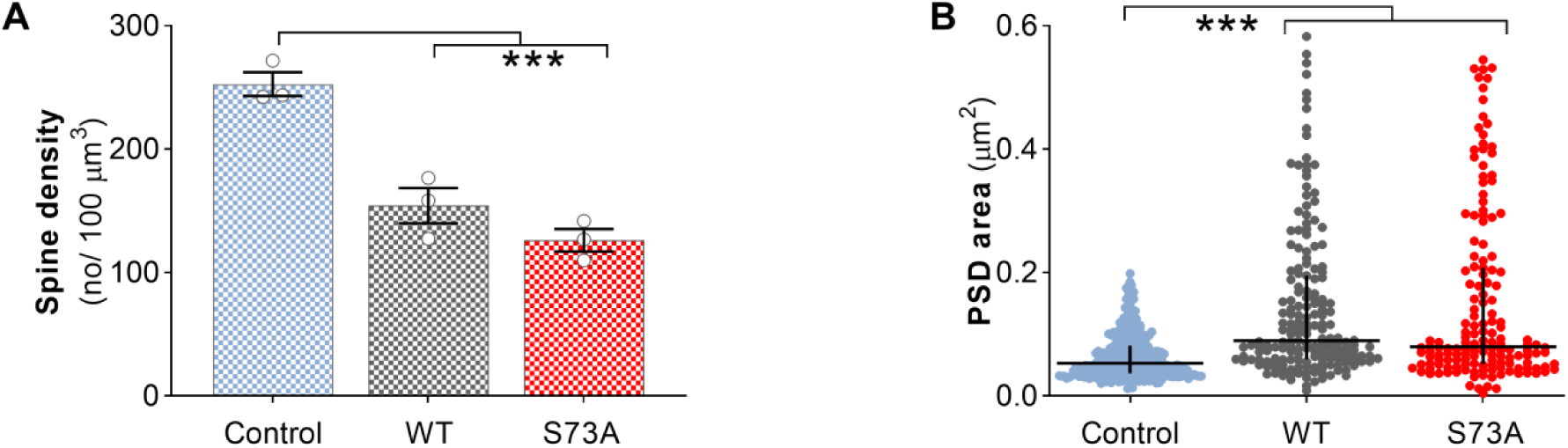
Overexpression of PSD-95 (WT and S73A) in dCA1 reduces dendritic spine density and increases PSDs size. Mice were stereotactically injected with AAVs encoding Control (mCherry), PSD-95 WT or S73A in the dCA1 and trained in contextual fear memory conditioning and extinction. (**A**) Mean density of dendritic spines was downregulated after overexpression of WT and S73A, compared to Control mice (One-way ANOVA, F(2, 6) = 34.59, ***P < 0.001). (**B**) Median size of PSDs was increased after overexpression of WT and S73A, compared to Control mice (Kruskal-Wallis statistic, H = 108.9, ***P < 0.001).

**Supplementary Figure 4.**
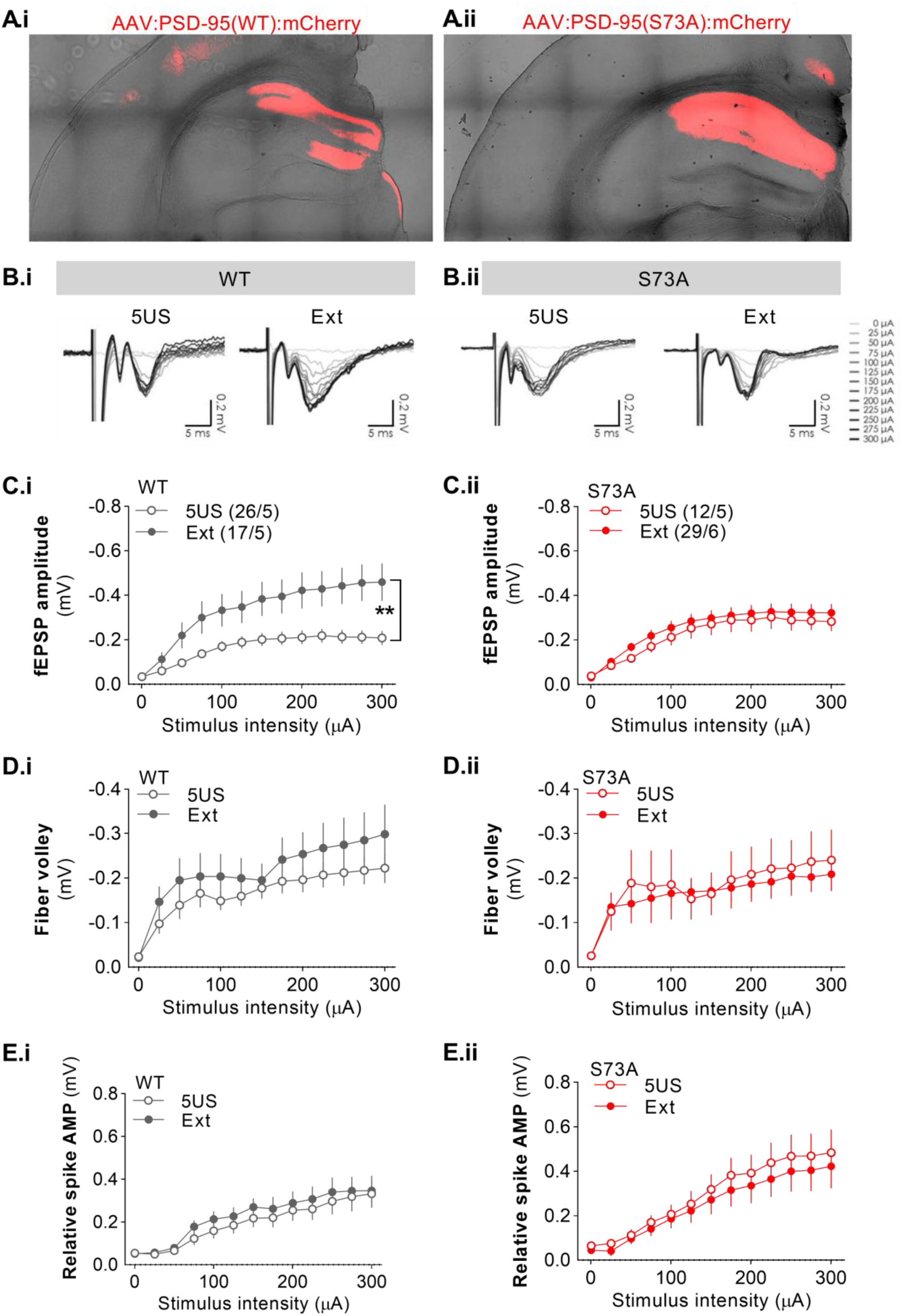
Phosphorylation of PSD-95 at S73 is required for fEPSP changes in stOri after fear extinction training. Mice were injected with PSD-95 WT or S73A in dCA1. Field excitatory postsynaptic potentials (fEPSPs) were recorded in stOri of dCA1 in response to the stimulation from the Schaffer collaterals. Moreover, the population spikes in the stratum pyramidale and fiber volleys were recorded and measured. **(A)** Microphotographs of PSD-95 WT and S73A expression in the CA1. **(B)** Representative fEPSPs evoked by stimuli of different intensities in stOri of the mice expressing PSD-95 WT and S73A respectively and sacrificed before or after fear extinction training. **(C)** Input–output plot for stimulus intensity versus fEPSP amplitude recorded in response to increasing intensities of stimulation in stOri. The fEPSP amplitudes were affected by the extinction training in the mice expressing **(i)** WT (RM ANOVA, F(1, 42) = 7.581, **P = 0.0087), **(ii)** but not S73A (F(1, 36) = 0.404, P = 0.528). (**D**) Input–output functions for stimulus intensity versus fiber volley recorded in response to increasing intensities of stimulation. No effect of contextual fear extinction was observed in mice expressing **(i)** WT (F(1, 30) = 0.080, P = 0.778) or **(ii)** S73A (F(1, 30) = 0.080, P = 0.778). The numbers of the analysed sections/mice per experimental group are indicated in (C).

**Supplementary Figure 5.**
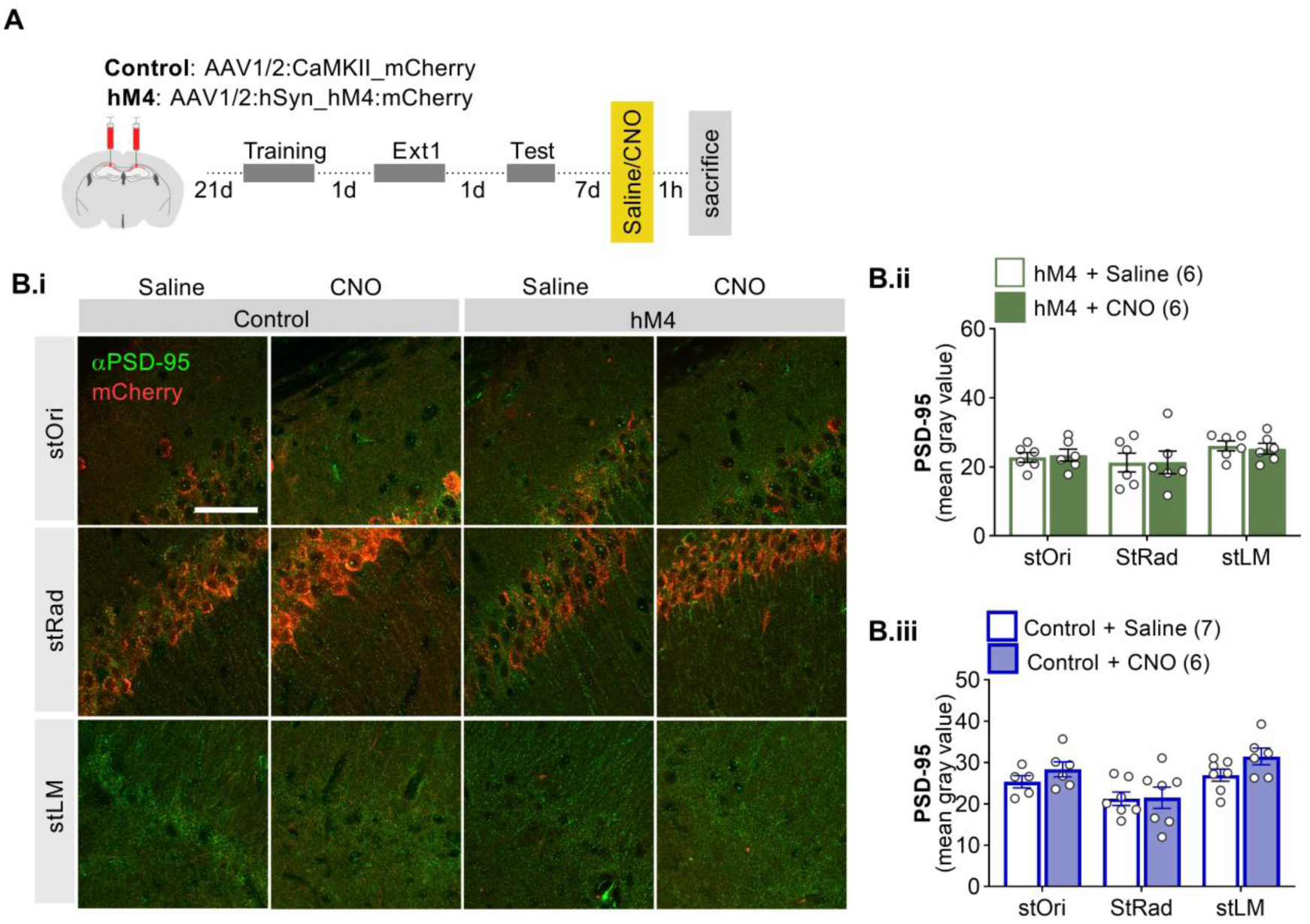
Chemogenetic inhibition of dCA1 after training does not affect PSD-95 expression. **(A)** Experimental timeline during fear conditioning and fear extinction sessions (ext1 and ext2) of the mice with Control and hM4 virus. Mice were trained three weeks after the surgery and virus expression. In all groups, saline or CNO (1 mg/kg) was systemically injected 7 days after the ext2. Mice were sacrificed 60 minutes after the injection. Behavioral data are shown in Fig. 2. **(B) (i)** Representative, confocal scans of the brain slices immunostained for PSD-95. Scale bar, 10 μm. (**ii-iii)** Summary of data quantifying the expression of PSD-95 in three domains of dCA1 in mice with Control or hM4 virus. No effect of the drug was observed in any of the virus groups (RM ANOVA, Control: F(1, 12) = 1.823, P = 0.322; hM4: F(1, 10) = 0.0003, P = 0.988). The numbers of mice per experimental group are in the legends (**B.ii-iii**).

## SUPPLEMENTARY MATERIALS AND METHODS

### Animals

C57BL/6J male mice were purchased from Białystok University, Poland. Thy1-GFP(M) (The Jackson Laboratory, JAX:007788, RRID:IMSR_JAX:007788) mutant mice were bred as heterozygotes at Nencki Institute, and PCR genotyped as previously described (Feng et al., 2000). All mice in the experiments were 7-9-week old. The mice were housed in groups of two to six and maintained on a 12 h light/dark cycle with food and water ad libitum. All experiments with transgenic mice used approximately equal numbers of males and females. The experiments were undertaken according to the Animal Protection Act of Poland and approved by the I Local Ethics Committee (261/2012, Warsaw, Poland).

### Contextual fear conditioning

The animals were trained in a conditioning chamber (Med Associates Inc, St Albans, USA) in a soundproof box. The chamber floor had a stainless steel grid for shock delivery. Before training, the chamber was cleaned with 70% ethanol, and a paper towel soaked in ethanol was placed under the grid floor. To camouflage background noise in the behavioral room, a white noise generator was placed inside the soundproof box.

On the conditioning day, the mice were brought from the housing room into a holding room to acclimatize for 30 min before training. Next, mice were placed in the training chamber, and after a 148 s introductory period, a foot shock (2 s, 0.7 mA) was presented. The shock was repeated 5 times, at 90 s inter-trial intervals. Thirty seconds after the last shock, the mouse was returned to its home cage. Contextual fear memory was tested and extinguished 24 h after training by re-exposing mice to the conditioning chamber for 30 minutes without US presentation, followed by the second 5-minute test session on the following day. During extensive contextual fear extinction, 30-minut fear extinction sessions were repeated on days 2, 3, 14, 15, and 16. Moreover mice activity and freezing were tested in context B (Ctx B) on day 17. A video camera was fixed inside the door of the sound attenuating box for the behavior to be recorded and scored. Freezing behavior (defined as complete lack of movement, except respiration) and locomotor activity of mice were automatically scored. The experimenters were blind to the experimental groups.

*CNO administration*. Clozapine N-Oxide (CNO) was dissolved in 0.9% saline. One or 3 mg/kg CNO was intraperitoneally (i.p.) injected 30 min before the behavioral extinction session. These doses of CNO did not induce any overt abnormal behaviors except for those reported in the study.

### Immunostaining

Mice were anesthetized and perfused with cold phosphate buffer pH 7.4, followed by 0.5% 4% PFA in phosphate buffer. Brains were removed and postfixed o/n in 4°C. Brains were kept in 30% sucrose in PBS for 72h. Coronal brain sections were prepared using cryosectioning (40 μm thick, Cryostat CM1950, Leica Biosystems Nussloch GmbH, Wetzlar, Germany) and stored in a cryoprotecting solution in –20°C (PBS, 15% sucrose (Sigma-Aldrich), 30% ethylene glycol (Sigma-Aldrich), and 0.05% NaN_3_ (SigmaAldrich). Before staining, sections were washed 3 × PBS and blocked for 1 hour at room temperature (RT) in 5% NDS with 0.3% Triton X-100 in PBS and then incubated o/n, 4°C with PSD-95 primary antibodies (1:500, Millipore, MAB1598, RRID:AB_11212185). On the second day slices were washed 3 × PBS with 0,3% Trition X-100 and incubated for 90 minutes with secondary antibodies conjugated with AlexaFluor 555 (1:500, Invitrogen, A31570, RRID:AB_2536180). Slices were then mounted on microscope slides (Thermo Fisher Scientific) and covered with coverslips in Fluoromount-G medium with DAPI (00-4959-52, Invitrogen).

### Confocal microscopy and image quantification

The microphotographs of dendritic spines in Thy1-GFP mice and fluorescent PSD-95 immunostaining were taken on a Spinning Disc confocal microscope (63 × oil objective, NA 1.4, pixel size 0.13 µm × 0.13 µm) (Zeiss, Göttingen, Germany). We took microphotographs (16 bit, z-stacks of 12-48 scans; 260 nm z-steps) of the dendrites from stratum oriens (stOri), stratum radiatum (stRad) and stratum lacunosum-moleculare (stLM) (6 dendrites per region per animal) of dorsal CA1 pyramidal neurons (AP, Bregma from -1.7 to 2.06). Each dendritic spine was manually outlined, and the spine area was measured with ImageJ 1.52n software measure tool. Custom-written Python scripts were used to analyze the mean gray value of PSD-95(+) puncta per dendritic spine.

The PSD-95 fluorescent immunostaining and PSD-95:mCherry over-expression were analyzed with Zeiss LSM 800 microscope equipped with Airy-Scan detection (63× oil objective and NA 1.4, pixel size 0.13 µm × 0.13 µm, 8 bit) (Zeiss, Göttingen, Germany). A series of 18 continuous optical sections (67.72 µm × 67.72 µm), at 0.26 μm intervals, were scanned along the z-axis of the tissue section. Six to eight z-stacks of microphotographs were taken per animal per region, from every sixth section through dCA1. Total PSD-95 levels was assessed as an image mean gray value. As shown by CLEM staining exogenous PSD-95:mCherry localises not only within dendritic spines but also dendrites (Figure 2E). However, the synaptic and dendritic PSD-95:mCherry puncta significantly differ in dimensions and intensity. Based on these differences synaptic PSD-95:mCherry was identified and analysed as roundish, small and very intensive puncta that clearly differ from the background (analysed in ImageJ with function Analyze Particles, with the size filter attribute set to: 0.00-0.70, and Threshold separately adjusted for each stratum by the experimenter blind to the experimental groups) (Figure 2E). Since dendritic PSD-95:mCherry is unlikely related to synaptic processes and PSD size it was ignored during the analysis (areas that were large and only slightly darker from the background). Exogenous synaptic PSD-95:mCherry levels were expressed as % area of ROI (Figure 2K).

### Stereotactic surgery

Mice were fixed in a stereotactic frame (51503, Stoelting, Wood Dale, IL, USA) and kept under isoflurane anesthesia (5% for induction, 1.5-2.0% during surgery). Adeno-associated viruses, serotype 1 and 2, (AAV1/2), solutions were injected into the dorsal CA1 area (Paxinos & Franklin 2001) at coordinates in relation to Bregma (AP, -2.1mm; ML, ±1.1 mm; DV, -1.3mm). 450 nl of AAV solutions were injected into the CA1 through a beveled 26 gauge metal needle, and 10 µl microsyringe (SGE010RNS, WPI, USA) connected to a pump (UMP3, WPI, Sarasota, USA), and its controller (Micro4, WPI, Sarasota, USA) at a rate 50 nl/ min. The needle was then left in place for 5 min, retracted +100 nm DV, and left for an additional 5 min to prevent unwanted spread of the AAV solution. Titers of AAV1/2 were: αCaMKII_PSD-95(WT):mCherry (PSD-95(WT)): 1.35 x10^9^/⌠l, αCaMKII_PSD-95(S73A):mCherry (PSD-95(S73A)): 9.12 x10^9^/⌠l), αCaMKII_mCherry (mCherry): viral titer 7.5 x10^7^/⌠l (obtained from Karl Deisseroth’s Lab), hSyn_HA-hM4D(Gi):mCherry (hM4) (Addgene plasmid #50475): 4.59 x10^7^/⌠l. Mice were allowed to recover from anesthesia for 2-3 h on a heating pad and then transferred to individual cages where they stayed until complete skin healing, and next, they were returned to the home cages. The viruses were prepared at the Nencki Institute core facility, Laboratory of Animal Models. After training, the animals were perfused with 4% PFA in PBS and brain sections from the dorsal hippocampus were immunostained for PSD-95 and imaged with Zeiss Spinning Disc confocal microscope (magnification: 10x) to assess the extent of the viral expression and PSD-95 expression.

### 3D electron microscopy

Mice were transcardially perfused with cold phosphate buffer pH 7.4, followed by 0.5% EM-grade glutaraldehyde (G5882 Sigma-Aldrich) with 2% PFA in phosphate buffer pH 7.4 and postfixed overnight in the same solution. Brains were then taken out of the fixative and cut on a vibratome (Leica VT 1200) into 100 μm slices. Slices were kept in phosphate buffer pH 7.4, with 0.1% sodium azide in 4°C for up to 14 days. For AAV-injected animals, the fluorescence of exogenous proteins was confirmed in all slices by fluorescent imaging. Then, slices were washed 3 × in cold phosphate buffer and postfixed with a solution of 2% osmium tetroxide (#75632 Sigma-Aldrich) and 1.5 % potassium ferrocyanide (P3289 Sigma-Aldrich) in 0.1 M phosphate buffer pH 7.4 for 60 min on ice. Next, samples were rinsed 5 × 3 min with double distilled water (ddH_2_O) and subsequently exposed to 1% aqueous thiocarbohydrazide (TCH) (#88535 Sigma) solution for 20 min. Samples were then washed 5 × 3 min with ddH_2_O and stained with osmium tetroxide (1% osmium tetroxide in ddH_2_O, without ferrocyanide) for 30 min in RT. Afterward, slices were rinsed 5 × 3 min with ddH_2_O and incubated in 1% aqueous solution of uranium acetate overnight in 4°C. The next day, slices were rinsed 5 × 3 min with ddH_2_O, incubated with lead aspartate solution (prepared by dissolving lead nitrate in L-aspartic acid as previously described (Deerinck et al., 2010)) for 30 min in 60°C and then washed 5 × 3 min with ddH_2_O and dehydration was performed using graded dilutions of ice-cold ethanol (30%, 50%, 70%, 80%, 90%, and 2 × 100% ethanol, 5 min each). Then slices were infiltrated with Durcupan resin. A(17 g), B(17 g) and D(0,51 g) components of Durcupan (#44610 Sigma-Aldrich) were first mixed on a magnetic stirrer for 30 min and then 8 drops of DMP-30 (#45348 Sigma) accelerator were added (Knott et al., 2009). Part of the resin was then mixed 1:1 (v/v) with 100% ethanol and slices were incubated in this 50% resin on a clock-like stirrer for 30 min in RT. The resin was then replaced with 100% Durcupan for 1 hour in RT and then 100% Durcupan infiltration was performed o/n with constant slow mixing. The next day, samples were infiltrated with freshly prepared resin (as described above) for another 2 hours in RT, and then embedded between flat Aclar sheets (Ted Pella #10501-10). Samples were put in a laboratory oven for at least 48 hours at65°C for the resin to polymerize. After the resin hardened, the Aclar layers were separated from the resin embedded samples, dCA1 region was cut out with a razorblade. Caution was taken for the piece to contain minimal resin. Squares of approximately 1 × 1 × 1 mm were attached to aluminium pins (Gatan metal rivets, Oxford instruments) with very little amount of cyanacrylamide glue. After the glue dried, samples were mounted to the ultramicrotome to cut 1 μm thick slices. Slices were transferred on a microscope slide, briefly stained with 1% toluidine blue in 5% borate and observed under a light microscope to confirm the region of interest (ROI). Next, samples were grounded with silver paint (Ted Pella, 16062-15) and pinned for drying for 4 – 12 hours, before the specimens were mounted into the 3View2 chamber.

### SBEM imaging and 3D reconstructions

Samples were imaged with Zeiss SigmaVP (Zeiss, Oberkochen, Germany) scanning electron microscope equipped with 3View2 chamber using a backscatter electron detector. Scans were taken in the middle portion of the CA1 stOri of the dorsal hippocampus. From each sample, 200 sections were collected (thickness 60 nm). Imaging settings: high vacuum with EHT 2.9-3.8 kV, aperture: 20 μm, pixel dwell time: 3 μs, pixel size: 5 – 6.2 nm. Scans were aligned using the ImageJ software (ImageJ - > Plugins -> Registration -> StackReg) and saved as .tiff image sequence. Next, alignment scans were imported to Reconstruct software (Fiala 2005), available at http://synapses.clm.utexas.edu/tools/reconstruct/reconstruct.stm (Synapse Web Reconstruct, RRID:SCR_002716). Spine density was analyzed from 3 bricks per animal with the unbiased brick method (Fiala and Harris 2001) per tissue volume. Brick dimensions 4.3 × 4.184 × 3 μm were chosen to exceed the length of the largest profiles in the data sets at least twice. To calculate the density of dendritic spines, the total volume of large tissue discontinuities was subtracted from the volume of the brick.

A structure was considered to be a dendritic spine when it was a definite protrusion from the dendrite, with electron-dense material (representing postsynaptic part of the synapse, PSD) on the part of the membrane that opposed an axonal bouton with at least 3 vesicles within a 50-nm distance from the cellular membrane facing the spine. For 3D reconstructions, PSDs and dendritic spines in one brick were reconstructed for each sample. PSDs were first reconstructed and second, their dendritic spines were outlined. To separate dendritic spine necks from the dendrites, a cut-off plane was used approximating where the dendritic surface would be without the dendritic spine. PSD volume was measured by outlining dark, electron-dense area on each PSD containing section. The PSD area was measured manually according to the Reconstruct manual. All non-synaptic protrusions were omitted in this analysis. For multi-synaptic spines, the PSD areas and volumes have been summed. In total, 1317 dendritic spines with their PSDs were manually segmented with the annotators blind to sample condition.

### Correlative light-electron microscopy (CLEM)

CLEM workflow was based on a previously established protocol with some modifications (Bishop et al., 2011). Mice infused with PSD-95(WT) in the CA1 were perfused as described above. Brains were then removed and postfixed o/n in 4°C. 100 µm thick brain slices were cut on a vibratome and embedded in low melting point agarose in phosphate buffer and mounted into imaging chambers. mCherry fluorescence in the stRad was photographed using Zeiss LSM800, z-stacks of 60 images (60 µm thick) at 63 × magnification. Next, the slice was transferred under the 2P microscope (Zeiss MP PA Setup), where a Chameleon laser was used to brand mark the ROI (laser length 870 nm, laser power 85%, 250 scans of each line). Then, SBEM staining was performed as described above. The resin-embedded hippocampus was then divided into 4 rectangles and each was mounted onto metal pins to locate the laser-induced marks. SBEM scanned within the laser marked frame. The fluorescent image was overlaid onto the SBEM image using dendrites and cell nuclei as landmarks using ImageJ 1.48k software (RRID:SCR_003070).

### Electrophysiology

Mice were deeply anesthesized with Isoflurane, decapitated and the brains were rapidly dissected and transfered into ice-cold cutting artificial cerebrospinal fluid (ACSF) consisting of (in mM): 87 NaCl, 2.5 KCl, 1.25 NaH2PO4, 25 NaHCO3, 0.5 CaCl2, 7 MgSO4, 20 D-glucose, 75 sacharose equilibrated with carbogen (5% CO2/95% O2). The brain was cut to two hemispheres and 350 μm thick coronal brain slices were cut in ice-cold cutting ACSF with Leica VT1000S vibratome. Slices were then incubated for 15 min in cutting ACSF at 32°C. Next the slices were transfered to recording ACSF containing (in mM): 125 NaCl, 2.5 KCl, 1.25 NaH_2_PO_4_, 25 NaHCO_3_, 2.5 CaCl_2_, 1.5 MgSO_4_, 20 D-glucose equilibrated with carbogen and incubated for minimum 1 hour at room temperature (RT).

Extracellular field potential recordings were recorded in a submerged chamber perfused with recording ACSF in RT. The potentials were evoked with a Stimulus Isolator (A.M.P.I Isoflex) with a concentric bipolar electrode (FHC, CBARC75) placed in the stOri of CA2 on the experiment. The stimulating pulses were delivered at 0.1 Hz and the pulse duration was 0.3 ms. Recording electrodes (resistance 1-4 MΩ) were pulled from borosilicate glass (WPI, 1B120F-4) with a micropipette puller (Sutter Instruments, P-1000) and filled with recording ACSF. The recording electrodes were placed in stOri of dCA1. Simultaneously, a second recording electrode was placed in the stratum pyramidale to measure population spikes. For each slice, the recordings were done in stOri. Recordings were acquired with MultiClamp 700B amplifier (Molecular Devices, California, USA), digitized with Digidata 1550B (Molecular Devices, California, USA) and pClamp 10.7 Clampex 10.0 software (Molecular Devices, California, USA). Input/output curves were obtained by increasing stimulation intensity by 25 μA in the range of 0-300 μA. All electrophysiological data was nalyzed with AxoGraph 1.7.4 software (Axon Instruments, U.S.A). The amplitude of fEPSP, relative amplitude of population spikes and fiber volley were measured.

### Statistics

Data are presented as mean ± standard error of the mean (SEM) for populations with normal distribution or as median ± interquartile range (IQR) for populations with non-normal distribution. An animal was used as a biological replication in all experiments except for the dendritic spine size distribution analysis. When the data met the assumptions of parametric statistical tests, results were analysed by one- or repeated measures two-way ANOVA, followed by Tukey’s or Fisher’s *post hoc* tests, where applicable. Data were tested for normality by using the Shapiro-Wilk test of normality and for homogeneity of variances by using the Levene’s test. For repeated-measure data with missing observation, a linear mixed model was used to analyze the results, followed by pairwise comparisons with Sidak adjustment for multiple comparisons. Areas of dendritic spines and PSDs did not follow normal distributions and were analysed with the Kruskal-Wallis test. Frequency distributions of PSD area to the spine volume ratio were compared with the Kolmogorov-Smirnov test. Correlations were analysed using Spearman correlation (Spearman r (s_r_) is shown), and the difference between slopes or elevation between linear regression lines was calculated with ANCOVA. Differences between the experimental groups were considered statistically significant if P < 0.05. Analyses were performed using the Graphpad Prism 8 or Statistica software. Mice were excluded from the analysis only if they did not express the tested virus in the target region, or the value exceeded 3 standard deviations from the mean.

## REFERENCES

Abraham WC, Jones OD, Glanzman DL. 2019. Is plasticity of synapses the mechanism of long-term memory storage? Npj Sci Learn 4:9. doi:10.1038/s41539-019-0048-y

Aziz W, Kraev I, Mizuno K, Kirby A, Fang T, Rupawala H, Kasbi K, Rothe S, Jozsa F, Rosenblum K, Stewart MG, Giese KP. 2019. Multi-input Synapses, but Not LTP-Strengthened Synapses, Correlate with Hippocampal Memory Storage in Aged Mice. Curr Biol 29:3600–3610.e4. doi:10.1016/j.cub.2019.08.064

Baldi E, Bucherelli C. 2015. Brain sites involved in fear memory reconsolidation and extinction of rodents. Neurosci Biobehav Rev 53:160–190. doi:10.1016/j.neubiorev.2015.04.003

Baldi E, Bucherelli C. 2014. Entorhinal cortex contribution to contextual fear conditioning extinction and reconsolidation in rats. Neurobiol Learn Mem 110:64–71. doi:10.1016/j.nlm.2014.02.004

Bannerman DM, Bus T, Taylor A, Sanderson DJ, Schwarz I, Jensen V, Hvalby Ø, Rawlins JNP, Seeburg PH, Sprengel R. 2012. Dissecting spatial knowledge from spatial choice by hippocampal NMDA receptor deletion. Nat Neurosci 15:1153–1159. doi:10.1038/nn.3166

Bannerman DM, Sprengel R, Sanderson DJ, McHugh SB, Rawlins JNP, Monyer H, Seeburg PH. 2014. Hippocampal synaptic plasticity, spatial memory and anxiety. Nat Rev Neurosci 15:181–192. doi:10.1038/nrn3677

Bats C, Groc L, Choquet D. 2007. The Interaction between Stargazin and PSD-95 Regulates AMPA Receptor Surface Trafficking. Neuron 53:719–734. doi:10.1016/j.neuron.2007.01.030

Bein O, Duncan K, Davachi L. 2020. Mnemonic prediction errors bias hippocampal states. Nat Commun 11:3451. doi:10.1038/s41467-020-17287-1

Béïque J, Andrade R. 2003. PSD-95 regulates synaptic transmission and plasticity in rat cerebral cortex. J Physiol 546:859–867. doi:10.1113/jphysiol.2002.031369

Berger-Sweeney J, Zearfoss NR, Richter JD. 2006. Reduced extinction of hippocampal-dependent memories in CPEB knockout mice. Learn Mem Cold Spring Harb N 13:4–7. doi:10.1101/lm.73706

Bevilaqua L, Bonini J, Rossato J, Izquierdo L, Cammarota M, Izquierdo I. 2006. The entorhinal cortex plays a role in extinction. Neurobiol Learn Mem 85:192–197. doi:10.1016/j.nlm.2005.10.001

Bitencourt RM, Pamplona FA, Takahashi RN. 2008. Facilitation of contextual fear memory extinction and anti-anxiogenic effects of AM404 and cannabidiol in conditioned rats. Eur Neuropsychopharmacol J Eur Coll Neuropsychopharmacol 18:849–859. doi:10.1016/j.euroneuro.2008.07.001

Bliss TVP, Collingridge GL. 1993. A synaptic model of memory: long-term potentiation in the hippocampus. Nature 361:31–39. doi:10.1038/361031a0

Cai C-Y, Chen C, Zhou Y, Han Z, Qin C, Cao B, Tao Y, Bian X-L, Lin Y-H, Chang L, Wu H-Y, Luo C-X, Zhu D-Y. 2018. PSD-95-nNOS Coupling Regulates Contextual Fear Extinction in the Dorsal CA3. Sci Rep 8:12775. doi:10.1038/s41598-018-30899-4

Chen X, Nelson CD, Li X, Winters CA, Azzam R, Sousa AA, Leapman RD, Gainer H, Sheng M, Reese TS. 2011. PSD-95 Is Required to Sustain the Molecular Organization of the Postsynaptic Density. J Neurosci 31:6329–6338. doi:10.1523/JNEUROSCI.5968-10.2011

Cheng D, Hoogenraad CC, Rush J, Ramm E, Schlager MA, Duong DM, Xu P, Wijayawardana SR, Hanfelt J, Nakagawa T, Sheng M, Peng J. 2006. Relative and absolute quantification of postsynaptic density proteome isolated from rat forebrain and cerebellum. Mol Cell Proteomics MCP 5:1158–1170. doi:10.1074/mcp.D500009-MCP200

Chetkovich DM, Bunn RC, Kuo S-H, Kawasaki Y, Kohwi M, Bredt DS. 2002. Postsynaptic targeting of alternative postsynaptic density-95 isoforms by distinct mechanisms. J Neurosci Off J Soc Neurosci 22:6415–6425. doi:20026598

de Oliveira Alvares L, Pasqualini Genro B, Diehl F, Molina VA, Quillfeldt JA. 2008. Opposite action of hippocampal CB1 receptors in memory reconsolidation and extinction. Neuroscience 154:1648–1655. doi:10.1016/j.neuroscience.2008.05.005

Denk W, Horstmann H. 2004. Serial Block-Face Scanning Electron Microscopy to Reconstruct Three-Dimensional Tissue Nanostructure. PLoS Biol 2. doi:10.1371/journal.pbio.0020329

Ehrlich I, Klein M, Rumpel S, Malinow R. 2007. PSD-95 is required for activity-driven synapse stabilization. Proc Natl Acad Sci 104:4176–4181. doi:10.1073/pnas.0609307104

Ehrlich I, Malinow R. 2004. Postsynaptic Density 95 controls AMPA Receptor Incorporation during Long-Term Potentiation and Experience-Driven Synaptic Plasticity. J Neurosci 24:916–927.

El-Boustani S, Ip JPK, Breton-Provencher V, Knott GW, Okuno H, Bito H, Sur M. 2018. Locally coordinated synaptic plasticity of visual cortex neurons in vivo. Science 360:1349–1354. doi:10.1126/science.aao0862

El-Husseini AE, Schnell E, Chetkovich DM, Nicoll RA, Bredt DS. 2000. PSD-95 involvement in maturation of excitatory synapses. Science 290:1364–1368.

Elkobi A, Ehrlich I, Belelovsky K, Barki-Harrington L, Rosenblum K. 2008. ERK-dependent PSD-95 induction in the gustatory cortex is necessary for taste learning, but not retrieval. Nat Neurosci 11:1149–1151. doi:10.1038/nn.2190

Feng G, Mellor RH, Bernstein M, Keller-Peck C, Nguyen QT, Wallace M, Nerbonne JM, Lichtman JW, Sanes JR. 2000. Imaging Neuronal Subsets in Transgenic Mice Expressing Multiple Spectral Variants of GFP. Neuron 28:41–51. doi:10.1016/S0896-6273(00)00084-2

Fischer A. 2004. Distinct Roles of Hippocampal De Novo Protein Synthesis and Actin Rearrangement in Extinction of Contextual Fear. J Neurosci 24:1962–1966. doi:10.1523/JNEUROSCI.5112-03.2004

Fitzgerald PJ, Pinard CR, Camp MC, Feyder M, Sah A, Bergstrom HC, Graybeal C, Liu Y, Schlüter OM, Grant SG, Singewald N, Xu W, Holmes A. 2015. Durable fear memories require PSD-95. Mol Psychiatry 20:901–912. doi:10.1038/mp.2014.161

Frankland PW, Bontempi B. 2005. The organization of recent and remote memories. Nat Rev Neurosci 6:119–130. doi:10.1038/nrn1607

Gardoni F, Polli F, Cattabeni F, Di Luca M. 2006. Calcium-calmodulin-dependent protein kinase II phosphorylation modulates PSD-95 binding to NMDA receptors. Eur J Neurosci 24:2694–2704. doi:10.1111/j.1460-9568.2006.05140.x

Garín-Aguilar ME, Díaz-Cintra S, Quirarte GL, Aguilar-Vázquez A, Medina AC, Prado-Alcalá RA. 2012. Extinction procedure induces pruning of dendritic spines in CA1 hippocampal field depending on strength of training in rats. Front Behav Neurosci 6. doi:10.3389/fnbeh.2012.00012

Gray JA. 1982. The neuropsychology of anxiety: An enquiry into the functions of the septo-hippocampal system. Behav Brain Sci 5:469–484. doi:10.1017/S0140525X00013066

Grossberg S, Merrill JW. 1992. A neural network model of adaptively timed reinforcement learning and hippocampal dynamics. Brain Res Cogn Brain Res 1:3–38. doi:10.1016/0926-6410(92)90003-a

Hirsch SJ, Regmi NL, Birnbaum SG, Greene RW. 2015. CA1-specific deletion of NMDA receptors induces abnormal renewal of a learned fear response. Hippocampus 25:1374–1379. doi:10.1002/hipo.22457

Hoover WB, Vertes RP. 2012. Collateral projections from nucleus reuniens of thalamus to hippocampus and medial prefrontal cortex in the rat: a single and double retrograde fluorescent labeling study. Brain Struct Funct 217:191–209. doi:10.1007/s00429-011-0345-6

Huh KH, Guzman YF, Tronson NC, Guedea AL, Gao C, Radulovic J. 2009. Hippocampal Erk mechanisms linking prediction error to fear extinction: Roles of shock expectancy and contextual aversive valence. Learn Mem 16:273–278. doi:10.1101/lm.1240109

Irvine EE, Vernon J, Giese KP. 2005. αCaMKII autophosphorylation contributes to rapid learning but is not necessary for memory. Nat Neurosci 8:411–412. doi:10.1038/nn1431

Ishizuka N, Weber J, Amaral DG. 1990. Organization of intrahippocampal projections originating from CA3 pyramidal cells in the rat. J Comp Neurol 295:580–623. doi:10.1002/cne.902950407

Ji J, Maren S. 2008. Differential roles for hippocampal areas CA1 and CA3 in the contextual encoding and retrieval of extinguished fear. Learn Mem 15:244–251. doi:10.1101/lm.794808

Kajiwara R, Wouterlood FG, Sah A, Boekel AJ, Baks-te Bulte LTG, Witter MP. 2008. Convergence of entorhinal and CA3 inputs onto pyramidal neurons and interneurons in hippocampal area CA1—An anatomical study in the rat. Hippocampus 18:266–280. doi:10.1002/hipo.20385

Kornau HC, Schenker LT, Kennedy MB, Seeburg PH. 1995. Domain interaction between NMDA receptor subunits and the postsynaptic density protein PSD-95. Science 269:1737–1740. doi:10.1126/science.7569905

Kumaran D, Maguire EA. 2006. An unexpected sequence of events: mismatch detection in the human hippocampus. PLoS Biol 4:e424. doi:10.1371/journal.pbio.0040424

Lattal KM, Abel T. 2004. Behavioral impairments caused by injections of the protein synthesis inhibitor anisomycin after contextual retrieval reverse with time. Proc Natl Acad Sci U S A 101:4667–4672. doi:10.1073/pnas.0306546101

Li J, Han Z, Cao B, Cai C-Y, Lin Y-H, Li F, Wu H-Y, Chang L, Luo C-X, Zhu D-Y. 2017. Disrupting nNOS-PSD-95 coupling in the hippocampal dentate gyrus promotes extinction memory retrieval. Biochem Biophys Res Commun 493:862–868. doi:10.1016/j.bbrc.2017.09.003

Lisman J, Buzsáki G, Eichenbaum H, Nadel L, Ranganath C, Redish AD. 2017. Viewpoints: how the hippocampus contributes to memory, navigation and cognition. Nat Neurosci 20:1434–1447. doi:10.1038/nn.4661

Mahmmoud RR, Sase S, Aher YD, Sase A, Gröger M, Mokhtar M, Höger H, Lubec G. 2015. Spatial and Working Memory Is Linked to Spine Density and Mushroom Spines. PLOS ONE 10:e0139739. doi:10.1371/journal.pone.0139739

Mamiya N, Fukushima H, Suzuki A, Matsuyama Z, Homma S, Frankland PW, Kida S. 2009. Brain Region-Specific Gene Expression Activation Required for Reconsolidation and Extinction of Contextual Fear Memory. J Neurosci 29:402–413. doi:10.1523/JNEUROSCI.4639-08.2009

Martin SJ, Grimwood PD, Morris RGM. 2000. Synaptic Plasticity and Memory: An Evaluation of the Hypothesis. Annu Rev Neurosci 23:649–711. doi:10.1146/annurev.neuro.23.1.649

Migaud M, Charlesworth P, Dempster M, Webster LC, Watabe AM, Makhinson M, He Y, Ramsay MF, Morris RGM, Morrison JH, O’Dell TJ, Grant SGN. 1998. Enhanced long-term potentiation and impaired learning in mice with mutant postsynaptic density-95 protein. Nature 396:433–439. doi:10.1038/24790

Morris RGM, Moser EI, Riedel G, Martin SJ, Sandin J, Day M, O’Carroll C. 2003. Elements of a neurobiological theory of the hippocampus: the role of activity-dependent synaptic plasticity in memory. Philos Trans R Soc Lond B Biol Sci 358:773–786. doi:10.1098/rstb.2002.1264

Nader K, Schafe GE, Le Doux JE. 2000. Fear memories require protein synthesis in the amygdala for reconsolidation after retrieval. Nature 406:722–726. doi:10.1038/35021052

Nagura H, Ishikawa Y, Kobayashi K, Takao K, Tanaka T, Nishikawa K, Tamura H, Shiosaka S, Suzuki H, Miyakawa T, Fujiyoshi Y, Doi T. 2012. Impaired synaptic clustering of postsynaptic density proteins and altered signal transmission in hippocampal neurons, and disrupted learning behavior in PDZ1 and PDZ2 ligand binding-deficient PSD-95 knockin mice. Mol Brain 5:43. doi:10.1186/1756-6606-5-43

Neves G, Cooke SF, Bliss TVP. 2008. Synaptic plasticity, memory and the hippocampus: a neural network approach to causality. Nat Rev Neurosci 9:65–75. doi:10.1038/nrn2303

Nikonenko I, Boda B, Steen S, Knott G, Welker E, Muller D. 2008. PSD-95 promotes synaptogenesis and multiinnervated spine formation through nitric oxide signaling. J Cell Biol 183:1115–1127. doi:10.1083/jcb.200805132

Nowacka A, Borczyk M, Salamian A, Wójtowicz T, Włodarczyk J, Radwanska K. 2020. PSD-95 Serine 73 phosphorylation is not required for induction of NMDA-LTD. Sci Rep 10:2054. doi:10.1038/s41598-020-58989-2

Pamplona FA, Bitencourt RM, Takahashi RN. 2008. Short- and long-term effects of cannabinoids on the extinction of contextual fear memory in rats. Neurobiol Learn Mem 90:290–293. doi:10.1016/j.nlm.2008.04.003

Ploghaus A, Tracey I, Clare S, Gati JS, Rawlins JN, Matthews PM. 2000. Learning about pain: the neural substrate of the prediction error for aversive events. Proc Natl Acad Sci U S A 97:9281–9286. doi:10.1073/pnas.160266497

Radulovic J, Tronson NC. 2010. Molecular Specificity of Multiple Hippocampal Processes Governing Fear Extinction. Rev Neurosci 21:1–18. doi:10.1515/REVNEURO.2010.21.1.1

Radwanska K, Medvedev NI, Pereira GS, Engmann O, Thiede N, Moraes MFD, Villers A, Irvine EE, Maunganidze NS, Pyza EM, Ris L, Szymańska M, Lipiński M, Kaczmarek L, Stewart MG, Giese KP. 2011. Mechanism for long-term memory formation when synaptic strengthening is impaired. Proc Natl Acad Sci U S A 108:18471–18475. doi:10.1073/pnas.1109680108

Ramanathan KR, Jin J, Giustino TF, Payne MR, Maren S. 2018. Prefrontal projections to the thalamic nucleus reuniens mediate fear extinction. Nat Commun 9:4527. doi:10.1038/s41467-018-06970-z

Restivo L, Vetere G, Bontempi B, Ammassari-Teule M. 2009. The Formation of Recent and Remote Memory Is Associated with Time-Dependent Formation of Dendritic Spines in the Hippocampus and Anterior Cingulate Cortex. J Neurosci 29:8206–8214. doi:10.1523/JNEUROSCI.0966-09.2009

Roth BL. 2016. DREADDs for Neuroscientists. Neuron 89:683–694. doi:10.1016/j.neuron.2016.01.040

Royer S, Paré D. 2003. Conservation of total synaptic weight through balanced synaptic depression and potentiation. Nature 422:518–522. doi:10.1038/nature01530

Sakaguchi M, Kim K, Yu LMY, Hashikawa Y, Sekine Y, Okumura Y, Kawano M, Hayashi M, Kumar D, Boyden ES, McHugh TJ, Hayashi Y. 2015. Inhibiting the Activity of CA1 Hippocampal Neurons Prevents the Recall of Contextual Fear Memory in Inducible ArchT Transgenic Mice. PLOS ONE 11.

Schafe GE, Nader K, Blair HT, LeDoux JE. 2001. Memory consolidation of Pavlovian fear conditioning: a cellular and molecular perspective. Trends Neurosci 24:540–546. doi:10.1016/S0166-2236(00)01969-X

Schnell E, Sizemore M, Karimzadegan S, Chen L, Bredt DS, Nicoll RA. 2002. Direct interactions between PSD-95 and stargazin control synaptic AMPA receptor number. Proc Natl Acad Sci U S A 99:13902–13907. doi:10.1073/pnas.172511199

Schuette PJ, Reis FMCV, Maesta-Pereira S, Chakerian M, Torossian A, Blair GJ, Wang W, Blair HT, Fanselow MS, Kao JC, Adhikari A. 2020. Long-Term Characterization of Hippocampal Remapping during Contextual Fear Acquisition and Extinction. J Neurosci 40:8329–8342. doi:10.1523/JNEUROSCI.1022-20.2020

Stansley BJ, Fisher NM, Gogliotti RG, Lindsley CW, Conn PJ, Niswender CM. 2018. Contextual Fear Extinction Induces Hippocampal Metaplasticity Mediated by Metabotropic Glutamate Receptor 5. Cereb Cortex 28:4291–4304. doi:10.1093/cercor/bhx282

Stein V, House DRC, Bredt DS, Nicoll RA. 2003. Postsynaptic Density-95 Mimics and Occludes Hippocampal Long-Term Potentiation and Enhances Long-Term Depression. J Neurosci 23:5503–5506. doi:10.1523/JNEUROSCI.23-13-05503.2003

Steiner P, Higley MJ, Xu W, Czervionke BL, Malenka RC, Sabatini BL. 2008. Destabilization of the Postsynaptic Density by PSD-95 Serine 73 Phosphorylation Inhibits Spine Growth and Synaptic Plasticity. Neuron 60:788–802. doi:10.1016/j.neuron.2008.10.014

Sturgill JF, Steiner P, Czervionke BL, Sabatini BL. 2009. Distinct Domains within PSD-95 Mediate Synaptic Incorporation, Stabilization, and Activity-Dependent Trafficking. J Neurosci 29:12845–12854. doi:10.1523/JNEUROSCI.1841-09.2009

Taft CE, Turrigiano GG. 2014. PSD-95 promotes the stabilization of young synaptic contacts. Philos Trans R Soc B Biol Sci 369:20130134. doi:10.1098/rstb.2013.0134

Tronson NC, Schrick C, Guzman YF, Huh KH, Srivastava DP, Penzes P, Guedea AL, Gao C, Radulovic J. 2009. Segregated Populations of Hippocampal Principal CA1 Neurons Mediating Conditioning and Extinction of Contextual Fear. J Neurosci 29:3387–3394. doi:10.1523/JNEUROSCI.5619-08.2009

Troyner F, Bertoglio LJ. 2021. Nucleus reuniens of the thalamus controls fear memory reconsolidation. Neurobiol Learn Mem 177:107343. doi:10.1016/j.nlm.2020.107343

Vallejo D, Codocedo JF, Inestrosa NC. 2017. Posttranslational Modifications Regulate the Postsynaptic Localization of PSD-95. Mol Neurobiol 54:1759–1776. doi:10.1007/s12035-016-9745-1

van Rooij SJH, Stevens JS, Ely TD, Hinrichs R, Michopoulos V, Winters SJ, Ogbonmwan YE, Shin J, Nugent NR, Hudak LA, Rothbaum BO, Ressler KJ, Jovanovic T. 2018. The Role of the Hippocampus in Predicting Future Posttraumatic Stress Disorder Symptoms in Recently Traumatized Civilians. Biol Psychiatry 84:106–115. doi:10.1016/j.biopsych.2017.09.005

Vertes RP, Linley SB, Hoover WB. 2015. Limbic circuitry of the midline thalamus. Neurosci Biobehav Rev 54:89–107. doi:10.1016/j.neubiorev.2015.01.014

Xu W, Sudhof TC. 2013. A Neural Circuit for Memory Specificity and Generalization. Science 339:1290–1295. doi:10.1126/science.1229534

Zanca RM, Sanay S, Avila JA, Rodriguez E, Shair HN, Serrano PA. 2019. Contextual fear memory modulates PSD95 phosphorylation, AMPAr subunits, PKMζ and PI3K differentially between adult and juvenile rats. Neurobiol Stress 10:100139. doi:10.1016/j.ynstr.2018.11.002.

